# A new role for Notch in the control of polarity and asymmetric cell division of developing T cells

**DOI:** 10.1101/628602

**Authors:** Mirren Charnley, Mandy Ludford-Menting, Kim Pham, Sarah M. Russell

## Abstract

A fundamental question in biology is how single cells can reliably produce progeny of different cell types. Notch signalling frequently facilitates fate determination. Asymmetric cell division (ACD) often controls segregation of Notch signalling by imposing unequal inheritance of regulators of Notch. Here, we assessed the functional relationship between Notch and ACD in mouse T cell development. To attain immunological specificity, developing T cells must pass through a pivotal stage termed β-selection, which involves Notch signalling and ACD. We assessed functional interactions between Notch and ACD during β-selection using direct presentation of Notch ligands, DL1 and DL4, and pharmacological inhibition of Notch signalling. Contrary to prevailing models, we find Notch controls distribution of Notch1 itself and cell fate determinants, α-Adaptin and Numb. Notch and CXCR4 signalling cooperated to drive polarity during division. Thus, Notch signalling directly orchestrates ACD, and Notch1 is differentially inherited by sibling cells.

## Introduction

Developing multicellular organisms must produce multiple cell types from one progenitor cell. This is achieved by sequential bifurcations in the fate of daughter cells following cell division. Fate bifurcation is driven by various means, including random differences in the daughters, different positional cues, or controlled segregation of fate determinants differentially into the two daughter cells, a process termed asymmetric cell division (ACD) (Dewey et al., 2015, Schweisguth, 2015, Venkei and Yamashita, 2018). A common mediator of fate bifurcation, which uses all of these processes, is the Notch signalling pathway (Artavanis-Tsakonas et al., 1999, Fortini, 2009, Liu et al., 2017, Sjoqvist and Andersson, 2017). Notch is a transmembrane receptor that responds to ligands belonging to the Delta and Jagged family, which are present on adjacent cells (Bray, 2016). Notch signalling directly regulates transcription to influence cell fate decisions, such as proliferation, death, multi-potency, differentiation and self-renewal (Fortini, 2009). Understanding how Notch signals differently in different tissue contexts is a subject of intense investigation (Bray, 2016). Differential Notch signalling in two daughter cells often occurs indirectly through the differential inheritance of regulators of Notch (Bray, 2016). Indeed, some of the most frequently observed fate determinants controlled by ACD are the Notch regulators, Numb and α-Adaptin (Daeden and Gonzalez-Gaitan, 2018, Dewey et al., 2015, Rhyu et al., 1994, Schweisguth, 2015). Consequently, the prevailing view is that, although an important executor of cell fate, Notch does not directly determine the direction of bifurcation in fate (Schweisguth, 2015, Venkei and Yamashita, 2018).

We considered the possibility that Notch might play a more active role in dictating the direction of fate bifurcation. One observation suggestive of a controlling role for Notch showed that Notch signalling was required for asymmetric distribution of Numb during ACD of *Drosophila* neural progenitors (Bhat, 2014, Pinto-Teixeira and Desplan, 2014). We have previously observed that Notch was polarised during interphase in developing T cells (Pham et al., 2015), leading to two non-mutually exclusive hypotheses: (i) that Notch might remain asymmetric during division, and thus be asymmetrically partitioned in the daughters, and (ii) that Notch might control partitioning of cell fate determinants during division. We tested those two hypotheses in this study using an *in vitro* model of T cell development.

T cell development is a highly tractable model for the study of cell fate, with well-established *in vitro* models and an extensive knowledge of the molecular pathways involved (Janas et al., 2010, Mohtashami et al., 2010, Schmitt and Zuniga-Pflucker, 2002, Van de Walle et al., 2013, Yui and Rothenberg, 2014). Many aspects of T cell biology share mechanisms of fate control with other developmental systems, and both Notch signalling and ACD operate in developing and mature T cells (Chang et al., 2007, Oliaro et al., 2010, Tajbakhsh et al., 2009). ACD occurs specifically during the β-selection phase of T cell development (Pham et al., 2015). In this critical phase, the developing T cells (termed DN3 cells) must balance self-renewal, genomic DNA recombination to produce a functional T cell receptor chain, and differentiation. Notch signalling is required for progression through β-selection and subsequent fate choices (Ciofani et al., 2004, Ciofani and Zuniga-Pflucker, 2005). We exploited two model systems to explore the role of Notch1 during division of developing T cells; the OP9-DL1 coculture, and surfaces functionalised with DL1 (Janas et al., 2010, Mohtashami et al., 2010, Schmitt and Zuniga-Pflucker, 2002, Van de Walle et al., 2013, Shukla et al., 2017). We show that the interaction of Notch1 with its ligands, DL1 and DL4, drives the polarisation of Notch1 itself, and the polarisation of cell fate determinants, α-Adaptin and Numb. Together, these data show that Notch signalling directly acts as a polarity cue, and that Notch1 is differentially inherited by the daughters of an ACD. This indicates that Notch plays a key role in dictating cell fate in developing T cells.

## Results

### Notch signalling drives polarisation during interphase and cell division

To discern whether Notch is a passive participant or driver of polarity in developing T cells, we first assessed the role of Notch signalling on the polarisation of developing T cells in interphase. Using established methods (Janas et al., 2010, Mohtashami et al., 2010, Schmitt and Zuniga-Pflucker, 2002, Van de Walle et al., 2013), we drove the development of mouse T cells from hematopoietic stem cells using the OP9-DL1 stromal cell line that expresses the Notch ligand, DL1. To ensure that the cells were poised to, but had not yet entered the β-selection checkpoint, we sorted for the DN3a stage by flow cytometry and reseeded onto OP9-DL1 cells. Notch signalling was disrupted using the γ-secretase inhibitor (dibenzazepine, DBZ) (De Kloe and De Strooper, 2014). As previously published (Ciofani et al., 2004), Notch inhibition in DN3 cells reduced differentiation to the DN4 and DP stages, reduced cell proliferation, and increased cell death (**Fig. S1**). To determine the role of Notch signalling in polarity, we performed immunofluorescence microscopy to assess the localisation of α-tubulin and markers of cell polarity on fixed DN3a cells in interphase. We chose an early time point (15 hours) where inhibiting Notch signalling had little or no impact on the number of attached DN3a cells (**Fig. 1A**), but slightly reduced microtubule organising centre (MTOC) polarisation to the interface between the DN3a cell and stromal cell (**Fig. 1B**). As previously seen (Pham et al., 2015), Notch1 was polarised to the interface **(Fig 1C)**. Remarkably, the polarisation of Notch1 was dependent upon its signalling capacity, as DBZ treatment substantially reduced Notch1 polarisation. CXCR4, the receptor for the chemokine CXCL12 which is required for progression through β-selection and regulates polarity in DN3 cells (Pham et al., 2015, Trampont et al., 2010), was also less polarised after DBZ treatment (**Fig. 1D**). The non-polarised control protein, CD25, was not affected by DBZ **(Fig. 1E)**. Two cell fate regulators that are polarised during ACD of developing T cells, α-Adaptin and Numb (Pham et al., 2015), also failed to polarise upon DBZ treatment (**Fig. 1F, G**). Thus, inhibiting Notch signalling reduced polarity of Notch1, CXCR4, α-Adaptin and Numb from above 70% to between 24 and 42%, similar to the level of polarisation observed with the CD25 control (Fig. 1E). These data suggest that Notch signalling was required for the polarisation of DN3a cells.

**Figure 1.**
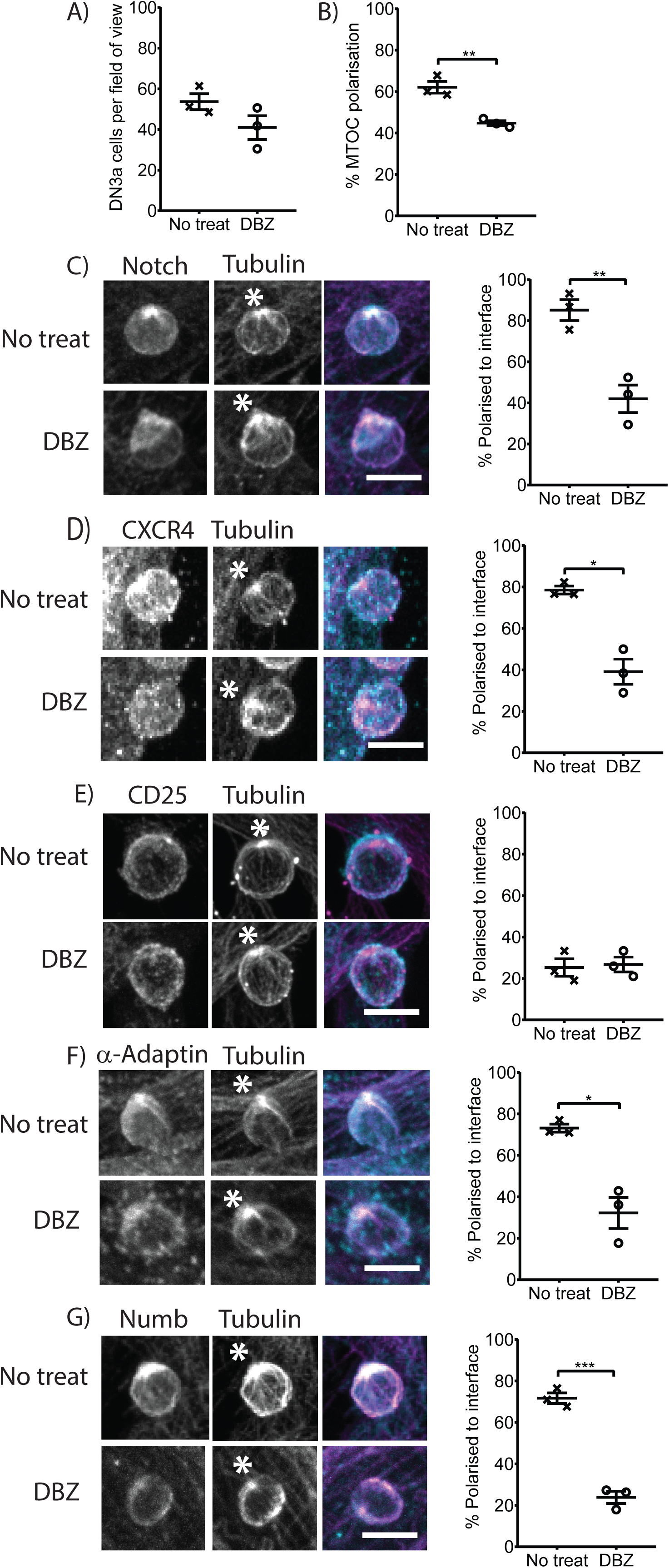
Intact Notch signalling is required for DN3a cell polarisation during interphase. DN3a cells were seeded on OP9-DL1 cells in the absence (‘no treat’) or presence of DBZ (0.05 µM) for 15 hours, fixed and stained for α-tubulin and Notch1, CXCR4, CD25, α-Adaptin or Numb. Inhibiting Notch signalling **(A)** did not affect thymocyte attachment to the stromal cell and **(B)** only slightly reduced MTOC recruitment to the interface between the stromal cell and the DN3a cell. **(C)** Notch1 and **(D)** CXCR4 were recruited to the cell-cell interface in over half the cells and this was severely disrupted in the presence of DBZ. **(E)** CD25 was not polarised and unperturbed by inhibiting Notch signalling. The cell fate regulators **(F)** α-Adaptin and **(G)** Numb were also polarised and this was reliant on intact Notch signalling. Representative projected z stack images are shown on the left, and quantification on the right. Total number of cells analysed: MTOC = 809 and 589, Notch1 = 130 and 110, CXCR4 = 158 and 113, CD25 = 68, 67, α-Adaptin = 131 and 105 and Numb = 158 and 110, for no treat and DBZ, respectively. Bar = 10 µm, * indicates the interface between the DN3a and OP9-DL1 stromal cells. *n* = 3 independent experiments. All data are represented as mean ± SEM; *p < 0.05, **p < 0.01, ***p < 0.001 (unpaired t test).

We next asked whether Notch1 was polarised during cell division, and if Notch signalling was required. Dividing cells were scored as undergoing symmetric cell division (SCD) if the protein of interest was symmetrically distributed between the two daughter cells, and as undergoing ACD if the protein of interest was more concentrated in one of the daughter cells. As previously seen (Pham et al., 2015), Numb and α-Adaptin were asymmetric in 55 to 66% of dividing cells, and we further show that Notch1 and CXCR4 were also asymmetric in 55 and 51% of dividing cells, while CD25 was asymmetric in only 29% **(Fig. 2A-E)**. Inhibiting Notch signalling reduced the asymmetry of the markers in dividing cells to the same level as the CD25 control. CXCR4 acts as a polarity cue for ACD (Pham et al., 2015), so the effect of Notch signalling on CXCR4 polarisation indicates that these cues could cooperate to trigger ACD. These data together indicate that Notch1 was polarised during division of DN3a cells, and suggest that Notch signalling is required for the polarisation of Notch1, CXCR4, α-Adaptin and Numb.

**Figure 2.**
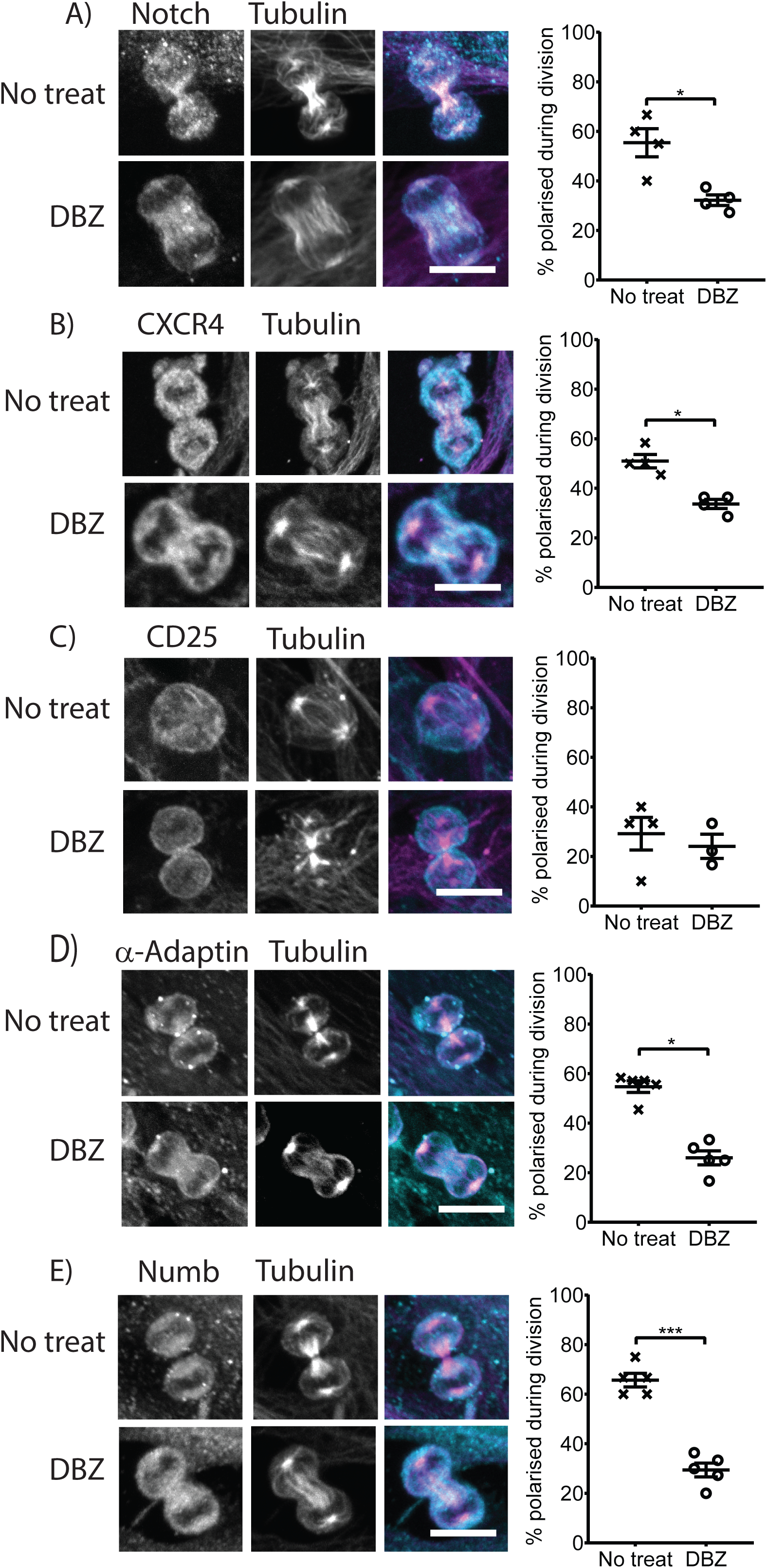
Intact Notch signalling is required for DN3a polarisation during cell division. DN3a cells were seeded on OP9-DL1 cells in the absence (‘no treat’) or presence of DBZ (0.05 µM) for 15 hours, fixed and stained for α-tubulin and either **(A)** Notch1, **(B)** CXCR4, **(C)** CD25, **(D)** α-Adaptin or **(E)** Numb. Representative projected z stack images are shown on the left, and quantification on the right. Inhibiting Notch signalling significantly reduced the percentage of cells undergoing ACD. Total number of cells analysed: Notch1 = 57 and 44, CXCR4 = 43 and 41, CD25 = 30 and 30, α-Adaptin = 46 and 56 and Numb = 40 and 45, for no treat and DBZ, respectively. Bar = 10 µm. *n* = 3-5 independent experiments. All data are represented as mean ± SEM; *p < 0.05, ***p < 0.001 (unpaired t test).

### Surfaces functionalised with the Notch ligand, DL1, were sufficient to induce polarisation during cell division

The pharmacological inhibition experiments above showed that γ-secretase was necessary for polarisation of DN3a cells during interphase and division. To confirm that these effects were through Notch, and to assess whether Notch signalling was sufficient for polarity, we used surfaces functionalised with the Notch ligand, DL1. DL1 was coupled to protein A (PA) coated surfaces via an Fc linker to control the orientation of DL1 relative to the cell (Makaraviciute and Ramanaviciene, 2013, Song et al., 2012, Toda et al., 2011). Growth factors essential to DN3a survival and differentiation were included in the cultures (Janas et al., 2010), but these were dispersed throughout the culture media and only the Notch ligand was presented to the cells with a defined orientation. As others have shown (Janas et al., 2010), surfaces functionalised with Fc-DL1 supported DN3 differentiation, albeit not as efficiently as OP9-DL1 cells (**Fig. S2**). The frequency and duration of divisions on the Fc-DL1 surfaces was comparable to that on the OP9-DL1 cells (**Fig. S3**). After two hours of culture the number of attached cells was unaffected by the surface coating but MTOC polarisation was increased on the Fc-DL1 surfaces, in comparison to PA alone (**Fig. 3A, B**). The recruitment of α-Adaptin and Numb to the interface was also increased on Fc-DL1 surfaces (**Fig. 3E-G**) and and this was maintained at 15 hours (Fig. 3C-G). These data demonstrate that the immobilised DL1 ligand alone was sufficient to trigger the recruitment of α-Adaptin and Numb to the functionalised surface.

**Figure 3.**
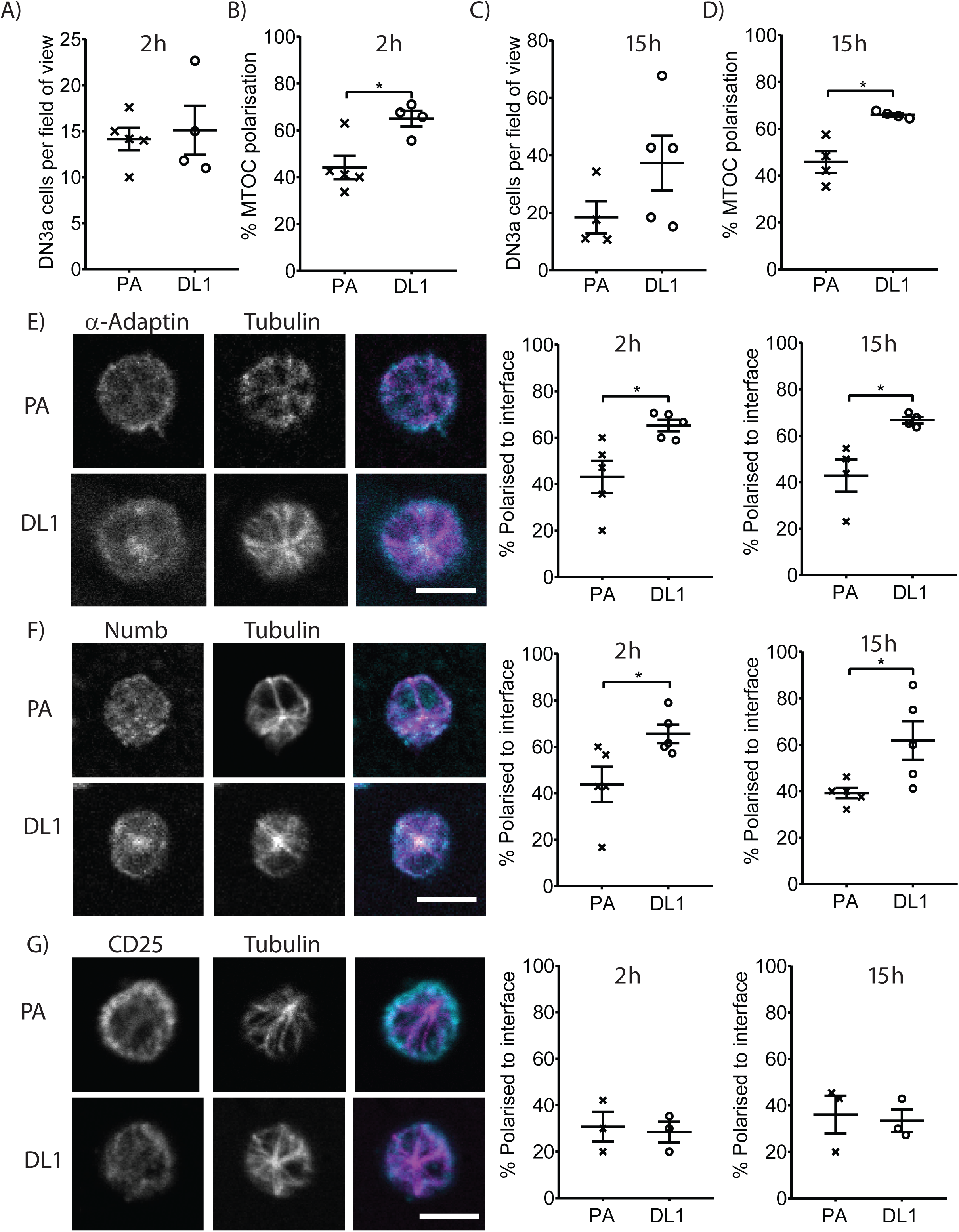
DN3a cells cultured on surfaces coated with the Notch ligand, DL1, polarise α-Adaptin and Numb during interphase. DN3a cells were cultured onto surfaces pre-coated with protein A (PA) or protein A and Fc-DL1 (DL1) for 2 or 15 hours before fixing. **(A)** After 2 hours thymocyte attachment was unaffected by the substrate coating, but **(B)** MTOC recruitment to the cell-substrate interface was increased in the presence of Fc-DL1 *versus* PA. A similar trend was seen after 15 hours **(C, D)**. Total number of cells analysed for 2 h = 115 (PA) and 122 (Fc-DL1) and 15 h = 77 (PA) and 157 (Fc-DL1). DN3a cells were cultured on surfaces pre-coated with PA or PA + Fc-DL1 (DL1) and stained with α-tubulin and either **(E)** α-Adaptin, **(F)** Numb or **(G)** CD25. Representative xy slice of the cell-substrate interface of DN3a cells on functionalised surfaces are shown on the left, and quantification on the right. α-Adaptin and Numb were recruited to the cell-cell interface on the Fc-DL1 functionalised substrates but not on the PA only substrates. Conversely, CD25 was not polarised regardless of the substrate. A similar trend in polarisation was observed at 2 and 15 h. Total number of cells analysed at 2 h: PA = 65, 74 and 39; DL1 = 54, 84 and 37 for α-Adaptin, Numb or CD25, respectively. Total number of cells analysed at 15 h: PA = 54, 69 and 35 and DL1 = 79, 124 and 38 for α-Adaptin, Numb or CD25, respectively. Bar = 10 µm. *n* = 3-5 independent experiments. All data are represented as mean ± SEM; *p < 0.05 (unpaired t test).

We then asked whether the surfaces coated with Fc-DL1 were capable of triggering asymmetry during division. First, DN3a cells were cultured on surfaces functionalised with Fc-DL1 or PA for 15 hours, fixed and labelled with α-tubulin and either α-Adaptin, Numb or CD25, imaged and blind scored using fluorescence microscopy. α-Adaptin and Numb had equal or greater polarity on Fc-DL1 functionalised surfaces, as compared to OP9-DL1 cells, and this response was specific to surfaces functionalised with DL1 (**Fig. 4A-C**). Similar to DN3a cells cultured on OP9-DL1 cells, Notch was asymmetric in 65% of dividing cells when cultured on Fc-DL1 functionalised surfaces and CXCR4 was asymmetric in 56% of cells (**Fig. 4D**). Given that both Notch1 and its regulators, α-Adaptin and Numb, were polarised, we assessed whether they were on the same or opposing sides of the dividing cell. For dividing cells in which both Notch1 and Numb were polarised, 84% showed recruitment of both proteins into the same daughter cell (**Fig. 4E**). Similarly, Notch1 was corecruited into the same daughter cell as α-Adaptin (86% of polarised divisions) and CXCR4 (79% of polarised divisions). This indicates that Notch ligation by DL1 drove polarisation during division, leading to the preferential recruitment of Notch1, CXCR4, Numb and α-Adaptin into one daughter cell.

**Figure 4.**
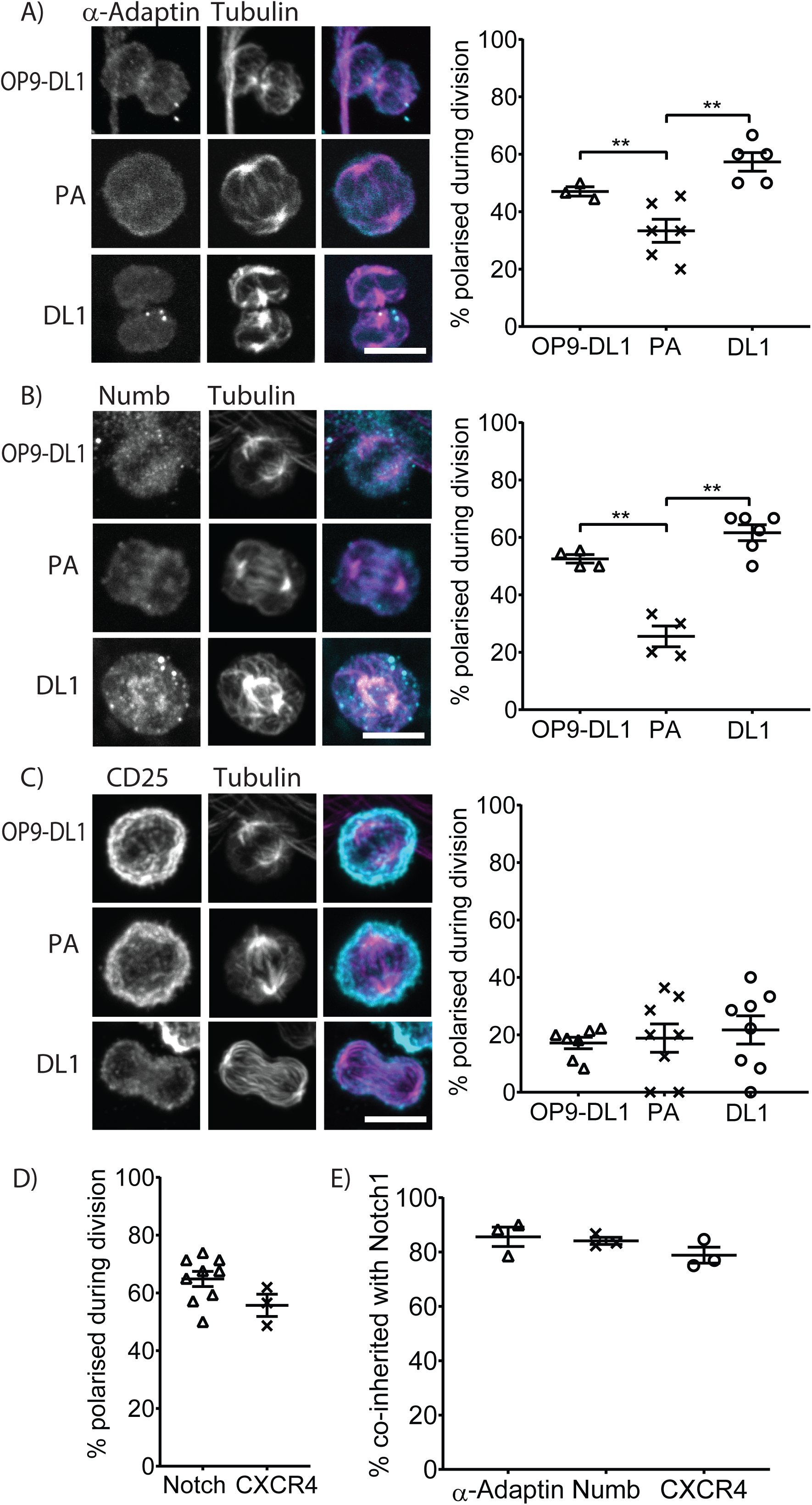
Dividing DN3a cells polarise cell fate determinants, α-Adaptin and Numb on functionalised surfaces. DN3a cells were cultured on surfaces pre-coated with PA or PA + Fc-DL1 (DL1) or OP9-DL1 stromal cells for 15 hours before fixing and staining with α-tubulin and either **(A)** α-Adaptin, **(B)** Numb or **(C)** CD25. Representative projected z stack images are shown on the left, and quantification on the right. α-Adaptin and Numb were asymmetrically distributed in a high proportion of cells when cultured on OP9-DL1 stromal cells or on Fc-DL1 functionalised surfaces, but not on surfaces only functionalised with PA. CD25 was symmetrically distributed regardless of the surface. Total number of divisions analysed: α-Adaptin = 40, 57 and 42; Numb = 46, 42 and 45; CD25 = 86, 73 and 67 for OP9-DL1, PA and Fc-DL1, respectively. To determine if Notch and the other cell fate determinanats were inherited into the same or opposing daughters cells, DN3a cells cultured on Fc-DL1 and stained with both Notch1 and the other cell fate determinants (i.e. α-Adaptin, Numb or CXCR4). **(D)** Notch and CXCR4 was asymmetrically distributed in a high proportion of divisions. **(E)** Dividing cells which were asymmetric for both proteins were selected for analysis and scored as co-inherited if both proteins were concentrated within the same nascent daughter cell. α-Adaptin, Numb and CXCR4 were co-inherited in the majority of dividing cells. Total number of divisions analysed: Notch = 244, CXCR4 = 81 (polarisation); α-Adaptin = 41, Numb = 44 and CXCR4 = 42 (co-inheritance with Notch1). All data are represented as mean ± SEM. **p < 0.01 (unpaired t test).

To complement the analysis of the fixed cells and quantify the extent of asymmetry, DN3a cells were transduced with fluorescently tagged α-Adaptin and Numb, cultured in cell paddocks onto which OP9-DL1 stromal cells had been allowed to attach overnight or were coated with PA or Fc-DL1 and imaged using time lapse microscopy (Day et al., 2009). The fluorescence intensity was quantified in dividing cells and expressed as polarisation ratios (PR; calculated as the difference in fluorescence between the two halves divided by the sum of the fluorescence. A low PR value indicates that the fluorescence is symmetric in the dividing cells, a high PR value indicates that it is asymmetric). A diffuse marker was also transduced into the DN3a cells (either GFP or cherry) to determine the background level of asymmetry and remove any artefacts (e.g. out of focus cells) (Charnley and Russell, 2017, Pham et al., 2015, Shimoni et al., 2014). The over-expression of the protein of interest had little or no effect on proliferation, differentiation or the level of polarity (**Fig. S4A-D**) (Pham et al., 2015, Pham et al., 2013). To ensure against possible artefacts related to the nature of the tag (Couturier et al., 2014, Short, 2014), we compared the polarity of α-Adaptin and Numb coupled to both GFP and cherry and saw no difference (**Fig. S4E, F**). Consequently, the data for the two fluorophores was combined (indicated as FP-α-Adaptin and FP-Numb). Dividing DN3a cells showed strong polarisation of both α-Adaptin (**Fig. 5A, S5A**) and Numb (**Fig. 5B, S5B**) on OP9-DL1 stromal cells. Dividing DN3a cells also showed strong polarisation of α-Adaptin and Numb when cultured on surfaces functionalised with DL1, which was comparable to that observed with OP9-DL1. Conversely, DN3a cells on the PA functionalised surfaces predominately divided symmetrically. As we previously observed with OP9-DL1 cells (Pham et al., 2015), asymmetric divisions occurred in DN3a cells but not in DN3b or DN2b cells (**Fig. S5C-E**). Thus, it is possible to mimic the stromal cell interaction and induce cell polarity by the orientated presentation of a single protein, namely Fc-DL1. This indicates that the binding of Notch1 to its ligand was both necessary and sufficient to dictate the polarisation of α-Adaptin and Numb during division.

**Figure 5.**
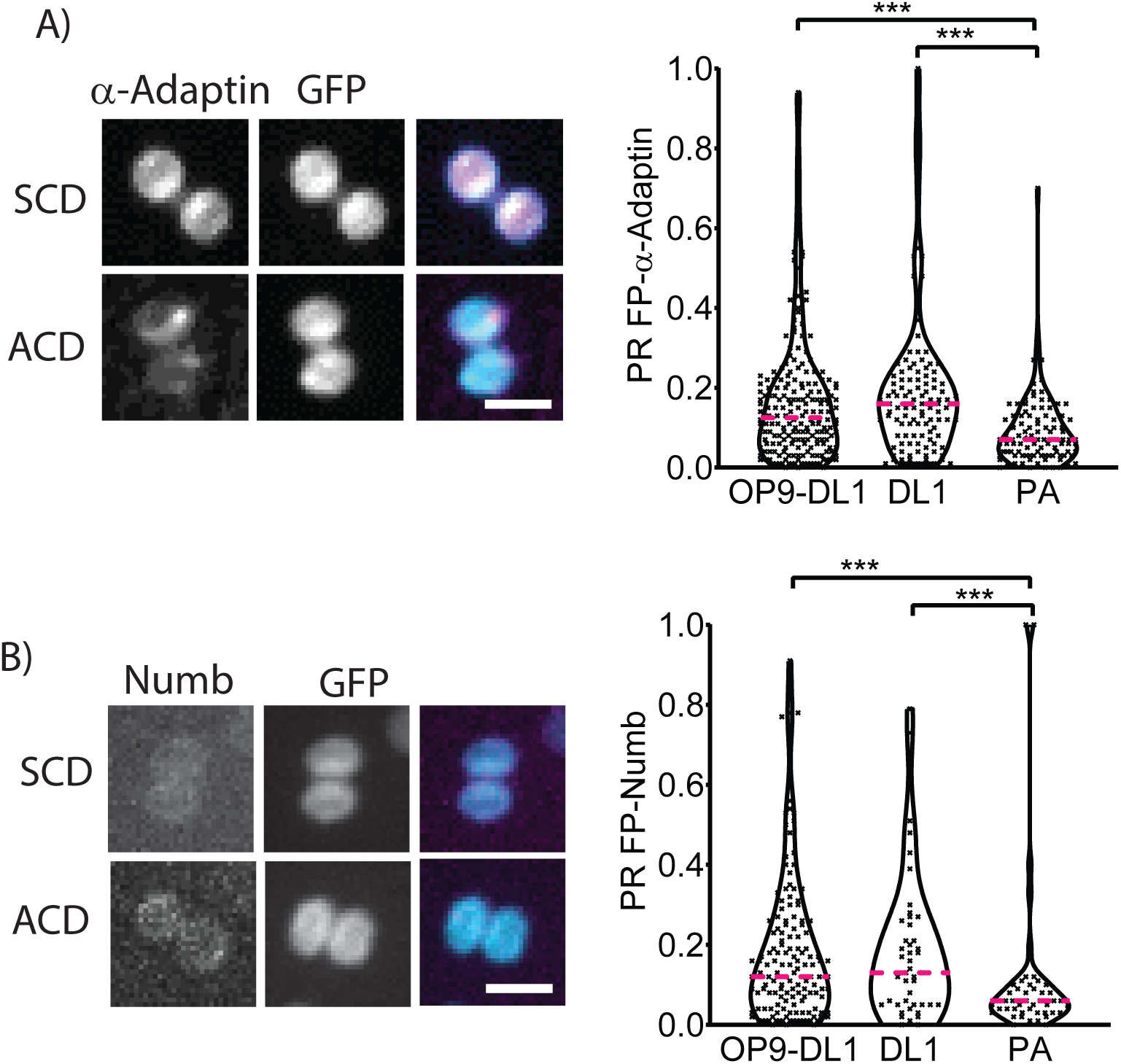
Dividing DN3a cells polarise cell fate determinants, α-Adaptin and Numb on functionalised surfaces. DN3a cells expressing a diffuse protein (GFP or cherry) and α-Adaptin were cultured onto surfaces pre-coated with PA, Fc-DL1 (DL1) or OP9-DL1 stromal cells and imaged using time-lapse microscopy for 20 hours. Representative projected z stack images are shown on the left, and quantification on the right. **(A)** Culturing DN3a cells on OP9-DL1 stromal cells and Fc-DL1 increased the proportion of dividing cells with high PRs. Number of divisions analysed: OP9-DL1 = 208, PA = 102 and Fc-DL1 = 109. **(B)** A similar trend was observed for DN3a cells transduced with Numb. Number of divisions analysed = OP9-DL1 = 148, PA = 48 and Fc-DL1 = 45. Bar = 10 µm. *n* = 3-8 independent experiments; magenta line on violin plots indicates median value. ***p < 0.001 (KS test). See Fig. S5 for PR scatter plots.

### Notch ligands presented on functionalised beads coordinate polarity and mitotic spindle orientation during ACD

ACD requires that polarity is coupled with orientation of the mitotic spindle (Venkei and Yamashita, 2018). We next asked whether the protein asymmetry triggered by Notch ligation was coordinated with the mitotic spindle. ACD of developing T cells requires that the mitotic spindle is orthogonal to the interface with the stromal cell (Pham et al., 2015). To determine the orientation of the cue relative to the axis of division, we used beads to pinpoint the localisation of the polarity cue (Habib et al., 2013), functionalised using the same strategy as the flat surfaces. The proportion of polarised divisions observed for the PA and Fc-DL1 functionalised beads was similar to the proportion observed for the flat surfaces (**Fig. 6A**). When the dividing cell was in contact with OP9-DL1 cells or Fc-DL1 functionalised beads, higher PRs correlated with greater division angles (**Fig. 6B, C**). Binning the data to separate cells based on division angle revealed that divisions perpendicular to the cue (division angles greater than 45°) showed higher PR values, indicating polarised divisions, compared with divisions that were parallel to the cue (**Fig. 6D, E**). This demonstrates that the protein asymmetry observed was coordinated with the orientation of the mitotic spindle, indicative of ACD. Thus, Notch ligands were necessary and sufficient to coordinate polarity and the spindle to orchestrate ACD.

**Figure 6.**
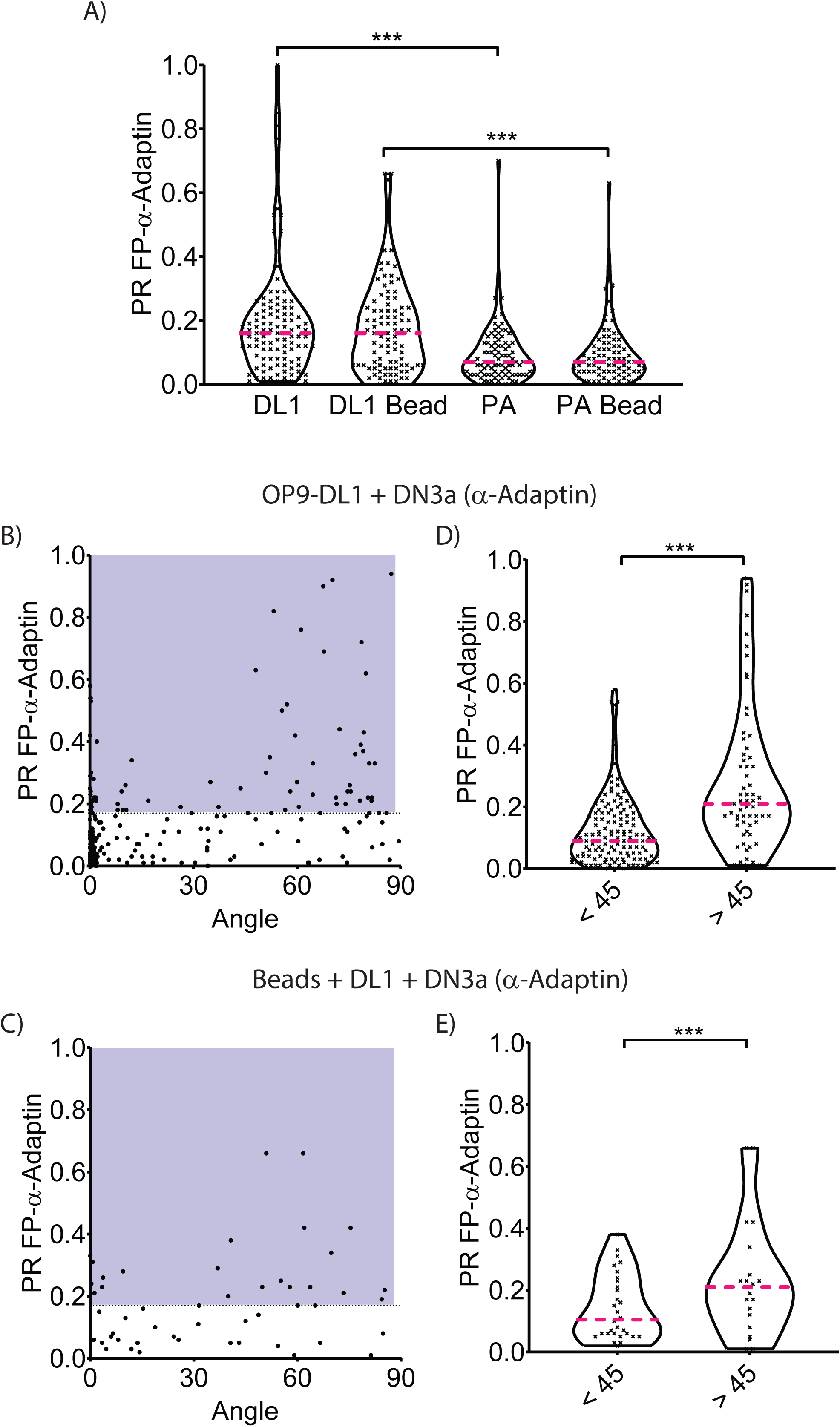
Polarity during division correlates with spindle orientation. DN3a cells expressing a diffuse protein (GFP or cherry) and test protein (α-Adaptin) were cultured in contact with 5 µm beads functionalised with Fc-DL1 (DL1) or OP9-DL1 stromal cells. **(A)** Beads functionalised with Fc-DL1, but not PA, triggered ACD, similar to functionalised flat surfaces (compare with Fig. 5). Total number of divisions analysed: DL1 = 109, DL1 bead = 94, PA = 102 and PA bead = 86. There was a correlation between increased division angles and polarity of α-Adaptin on **(B)** OP9-DL1 stromal cells or **(C)** Fc-DL1 functionalised beads. The blue shaded region indicates divisions that were assigned as asymmetric when the 0.17 cut-off was applied. **(D, E)** The same trend was observed when we compared the spread in PR values for cells which divided with a division angle < 45° *versus* > 45°. Number of divisions analysed: OP9-DL1 = 200; Fc-DL1 beads = 51. *n* = 3-8 independent experiments; magenta line on violin plots indicates median value; ***p < 0.001 (KS test).

### Notch functionally interacts with other cues to orchestrate ACD

The only directional cue presented to DN3a cells that promoted polarisation in the above experiments was DL1. However, the CXCR4 ligand, CXCL12, is required for T cell differentiation (Janas et al., 2010, Tussiwand et al., 2011) and was included in our cultures. We have previously found that ACD induced by OP9-DL1 stromal cells depends upon CXCR4 signalling in DN3a cells (Pham et al., 2015). This, combined with our observations above that CXCR4 polarisation depends on Notch signalling and that surfaces functionalised with the Notch ligand, DL1, induce asymmetry of α-Adaptin and Numb in dividing DN3a cells, raises the question of how these two receptors cooperate. We therefore assessed whether cells cultured on Fc-DL1 functionalised surfaces could still polarise after pharmacological inhibition of one or both of the CXCR4 and Notch signalling pathways, using AMD3100 or DBZ, respectively (**Fig. 7A, S6A-C**). Inhibiting either pathway resulted in a decrease in the polarisation of α-Adaptin. Further, both Notch and CXCR4 signalling was required for optimal polarisation of α-Adaptin in dividing DN3a cells, with the least polarisation occurring when both inhibitors were present. The effect of disrupting CXCR4 signalling is intriguing given that CXCL12 is diffuse in the cultures, so we looked specifically at the localisation of the two receptors. As expected, Notch1 was polarised in response to surface-bound DL1 in dividing DN3a cells (**Fig. 7B**). Notch1 polarisation was strongly inhibited by the Notch inhibitor, but only minimally affected by inhibition of CXCR4 signalling. CXCR4 was also polarised in response to presentation of the Notch ligand, and this polarisation was dependent on signalling through both Notch1 and CXCR4 (**Fig. 7C**). These data indicate that Notch ligation can drive the asymmetric polarisation of CXCR4 and that CXCR4 and Notch cooperate as non-redundant cues for ACD, albeit with Notch playing a more dominant role. Collectively these data show that multiple cues can interact to enhance or reduce the level of ACD.

**Figure 7.**
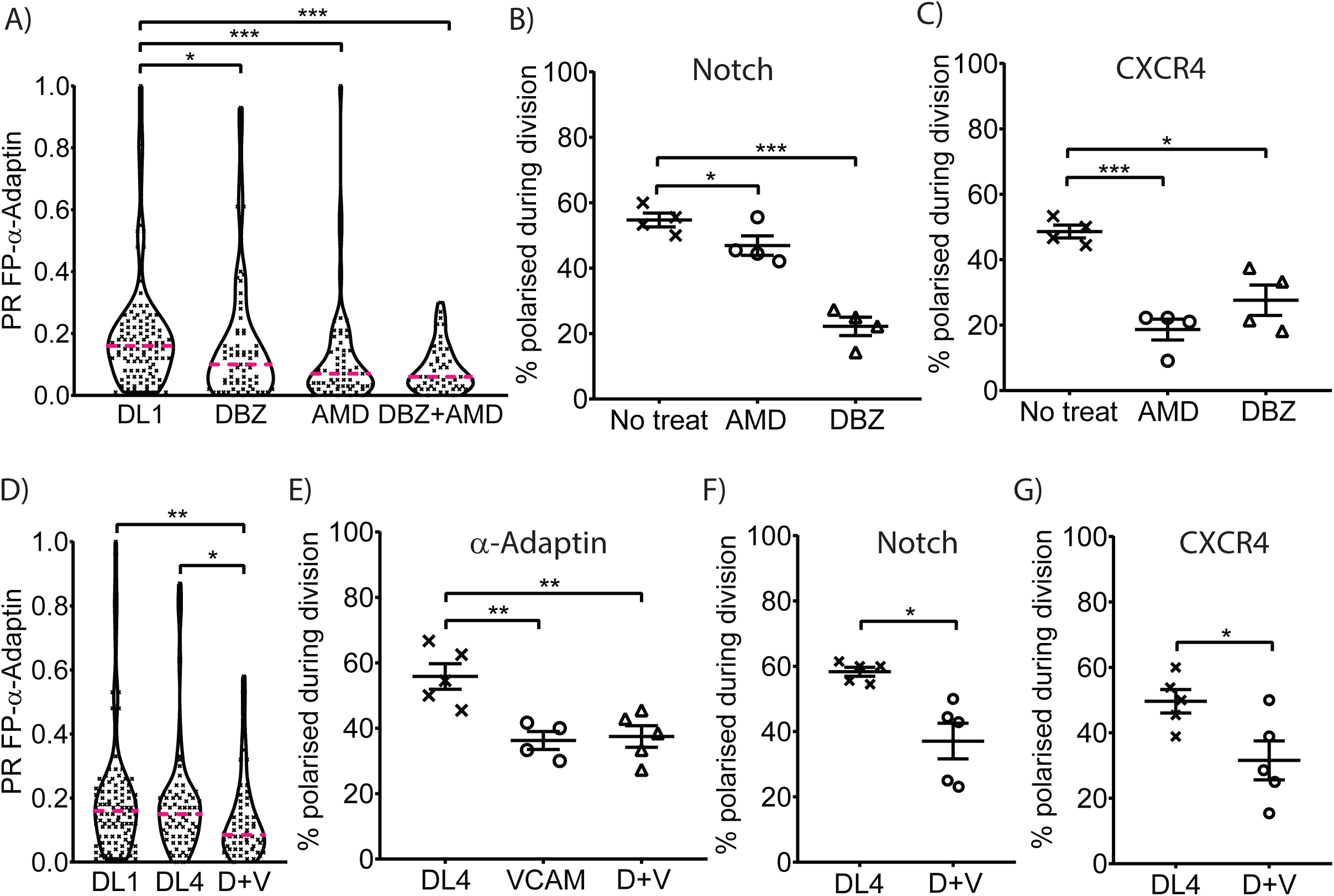
Different polarity cues cooperate to regulate ACD. DN3a cells expressing a diffuse protein (GFP or cherry) and α-Adaptin were cultured onto surfaces pre-coated with PA, Fc-DL1 (DL1), Fc-DL4 (DL4) or Fc-DL4 + Fc-VCAM-1 (D+V) and imaged using time-lapse microscopy for 20 hours. The addition of **(A)** either DBZ or AMD3100 (AMD) or a combination of both inhibitors caused a reduction in the level of ACD. Number of divisions: DBZ = 68, AMD3100 = 65, DBZ + AMD3100 = 51. **(B)** The polarisation of Notch1 in dividing cells was reduced by DBZ, but only slightly effected by AMD3100. **(C)** Conversely, CXCR4 polarisation was reduced by the inhibition of either Notch or CXCR4 signalling. Number of divisions analysed = 55, 48 and 42 for no treat, DBZ and AMD3100, respectively. **(D)** Fc-DL4 coated surfaces triggered ACD and the inclusion of Fc-VCAM-1 lead to a reduction in the asymmetric segregation of α-Adaptin during division. Number of divisions analysed: DL4 = 67; DL4 + VCAM = 58. The presence of VCAM-1 in the functionalised surfaces also resulted in reduced polarisation of **(E)** α-Adaptin, **(F)** Notch1 and **(G)** CXCR4 in dividing cells. Number of divisions analysed: DL4 = 56; DL4 + VCAM = 44 and DL4 = 64 (α-Adaptin); DL4 = 67; DL4 + VCAM = 71 (Notch1 and CXCR4). n = 3-6 independent experiments; scatter plots represent mean ± SEM, magenta line on violin plots indicates median value; * p < 0.05, **p < 0.01, ***p < 0.001 (KS test for violin plots and unpaired t test for scatter plots). See Fig. S6 for PR scatter plots.

Given the complexity of interactions in the thymus (Petrie and Zúñiga-Pflücker, 2007), we evaluated two other ligands for the regulation of ACD in DN3a cells. DL4 is an alternative ligand for Notch1 (Koch et al., 2008, Hozumi et al., 2008, Hozumi et al., 2004), and we assessed whether it behaved similarly to DL1. Indeed, both ligands induced equivalent polarisation of α-Adaptin in dividing DN3a cells (**Fig. 7D, S6D**). VCAM-1 contributes to stromal-DN3 interactions and its presence is associated with improved i*n vitro* generation of T cells (Shukla et al., 2017, Abe et al., 2010). We therefore assessed the impact of Fc-VCAM-1 on DL4-mediated polarisation during division by functionalising the surfaces with both ligands. Interestingly, the inclusion of Fc-VCAM-1 slightly reduced the level of polarisation (**Fig. 7D, E, S6E**). One possible explanation for this decrease is that the presence of VCAM-1 altered the interaction of the DL1 and CXCL12 ligands with their receptors. Indeed, on the Fc-DL4 + Fc-VCAM-1 surfaces the polarisation of both Notch1 and CXCR4 was also slightly reduced (**Fig. 7F, G**). This effect correlated with a decrease in polarisation at interphase of the MTOC and α-Adaptin (**Fig. S7A-C**), but was not associated with an effect on differentiation (**Fig. S7D-F**). These data show that the slight reduction in ACD observed when VCAM-1 was included in the functionalised surfaces was associated with reduced polarisation of Notch1 and CXCR4.

## Discussion

It is well established that Notch controls diversification in cell fate in a number of developmental systems. However, most of the proposed mechanisms assume that Notch executes decisions that have been made stochastically or are imposed by unequal expression of regulators of Notch (Dewey et al., 2015, Schweisguth, 2015, Venkei and Yamashita, 2018). Here, we demonstrate that Notch1 can directly trigger asymmetry in daughter cells, by steering ACD and by partitioning itself asymmetrically in the two daughter cells (**Fig. 8)**.

**Figure 8.**
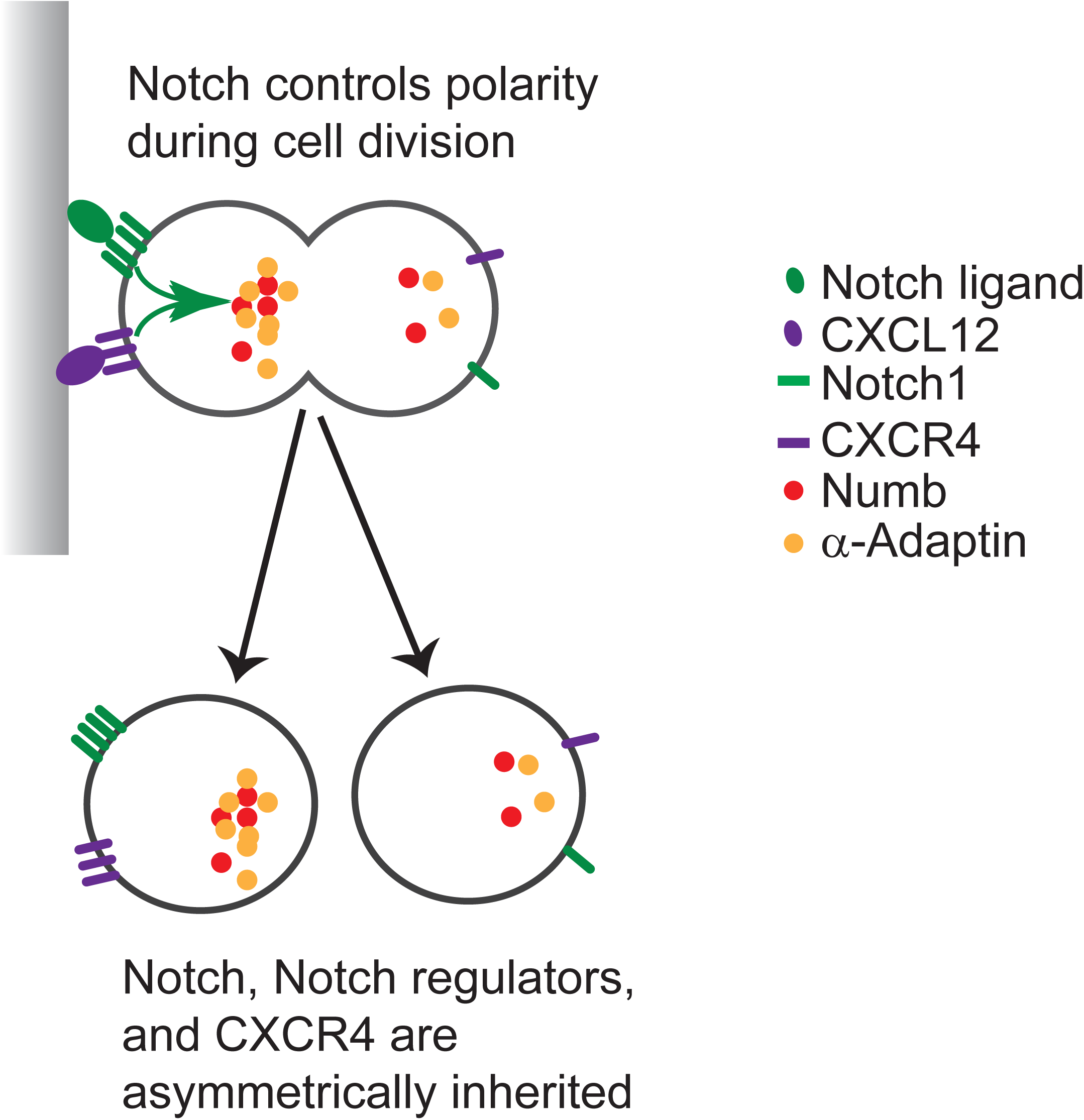
A model of the role of Notch in controlling cell polarity and ACD. Notch ligation of DL1, presented either by the OP9-DL1 stromal cell or DL1 functionalised surfaces, orchestrates polarisation during the division of DN3a cells. CXCR4 signalling cooperates with Notch to drive this asymmetry during division. This results in the differential inheritance of the cell fate determinants, α-Adaptin and Numb, CXCR4 and Notch1 itself, which were preferentially recruited into the same daughter cell.

Two model systems were used to explore the role of Notch1 during division of developing T cells; the OP9-DL1 coculture, and surfaces functionalised with Fc-DL1. The OP9-DL1 coculture system has been extensively used to study T cell development *in vitro*, and mimics physiological early T cell development with high fidelity (Janas et al., 2010, Mohtashami et al., 2010, Schmitt and Zuniga-Pflucker, 2002, Van de Walle et al., 2013). Surfaces functionalised with Notch ligands have also provided insights into the mechanisms controlling thymocyte development (Janas et al., 2010, Tussiwand et al., 2011, Dallas et al., 2005). Further, there is a growing interest in using surfaces functionalised with Notch ligands for the *in vitro* generation of T cells for therapeutic treatments, such as cancer immunology ((Gehre et al., 2015, Shukla et al., 2017, Suzuki et al., 2006) and as reviewed in (Singh and Zuniga-Pflucker, 2018)). Recently it was demonstrated that it is possible to direct human hematopoietic stem cells along the T cell lineage utilising functionalised surfaces in serum-free conditions (Shukla et al., 2017), which would circumvent issues associated with xenogeneic culture and enable the generation of clinical-grade pro-T cells.

Functionalised surfaces for the controlled presentation of proteins have been widely used to study fundamental biology, including adhesion and migration (Charnley et al., 2012, Gavard et al., 2004, Kovacs et al., 2002, Silvestre et al., 2009), division (den Elzen et al., 2009, Toyoshima and Nishida, 2007, Charnley et al., 2013), immune responses (Grakoui et al., 1999, Manz and Groves, 2010, Mossman et al., 2005, Charnley et al., 2009) and T cell development (Janas et al., 2010, Tussiwand et al., 2011, Dallas et al., 2005). Here, we demonstrated that these surfaces successfully mimicked another aspect of T cell development; developing T cells cultured on the surfaces underwent ACD and this process was coordinated by external cues and restricted to the DN3a stage of development. This work also highlights that functionalised surfaces can be used to manipulate the cells’ microenvironment and systemically test the influence of extrinsic factors on ACD.

The interaction of Notch1 with the DL1 ligand drove the polarisation of fate determinants that are asymmetrically distributed in several models of ACD, namely α-Adaptin and Numb. Notch signalling also triggered the polarisation of CXCR4, which is a polarity cue for ACD of DN3a cells (Pham et al., 2015), and the polarisation of Notch1 itself. Thus, the orientated presentation of the Notch1 ligand was sufficient to trigger polarity during division in developing T cells. The role of Notch in controlling localisation of Numb is analogous to that observed in *Drosophila* ganglion mother cells (Bhat, 2014), indicating that its role as a polarity cue could be more broadly conserved.

In addition to its role in controlling polarity, we demonstrate that, in dividing DN3a cells, Notch1 was asymmetrically distributed between the two daughter cells. Asymmetric partitioning of Notch1 itself during an ACD provides direct mechanism to establish variation in Notch signalling levels. Interestingly, Notch1 localisation is also controlled at cytokinesis in dividing sensory organ precursor cells (Coumailleau et al., 2009, Couturier et al., 2012, Trylinski et al., 2017). However, in these cells Notch was localised at the interface between two daughter cells during division, rather than polarised to one side of the mother cell. Using developing T cells as a model system of ACD we demonstrated that Notch1 is asymmetrically inherited, which could provide a direct and novel mechanism to establish differential Notch signalling in the two daughters (Fig 8).

More work is needed to clarify how this impacts on cell fate determination during β-selection. Notch1 signalling is required for T cell development past the β-selection stage, but peaks at the DN3a stage (Mingueneau et al., 2013) and is dramatically downregulated following β-selection (Yashiro-Ohtani et al., 2009). The role of Notch1 in enhancing ACD, and our finding that ACD is restricted to DN3a cells (current study and (Pham et al., 2015)) suggests the possibility that Notch acts in part to determine the developmental stage at which ACD occurs. Additionally, the segregation of Notch signalling by ACD might enable one daughter cell to progress past the β-selection stage, and the other to self-renew. The coinheritance of Notch antagonists with the Notch receptor suggests that this daughter cell is primed, but not necessarily active, for Notch signalling. In contrast, the daughter cell that receives less Notch1 presumably has less need to express the Notch antagonists. However, the endurance of this balance of inheritance of positive and negative regulators of Notch signalling, and its effect on cell fate will require analysis of the dynamics of Notch signalling over several generations.

Notch1, pre-TCR and CXCR4 signalling cooperate to drive the developing T cell through β-selection, and the presence of both CXCR4 and Notch1 signalling is required for survival on the functionalised surfaces (Ciofani et al., 2004, Ciofani and Zuniga-Pflucker, 2005, Janas et al., 2010, Trampont et al., 2010). Our results indicate that they also cooperate to dictate polarity and ACD in developing T cells. A functional interaction between Notch1 and CXCR4 has been demonstrated in a number of cell systems (Chiaramonte et al., 2015, Cong et al., 2017, Ho et al., 2017, Wang et al., 2009, Xiao et al., 2017), but the molecular basis for this interaction is not clear. The co-recruitment of Notch1 and CXCR4 could indicate that they physically interact, either via a direct interaction of the receptors or their ligands. The striking loss of CXCR4 polarisation when Notch signalling is inhibited suggests that Notch is upstream of the chemokine receptor. Taken together, these findings indicate that Notch1 and CXCR4 have complementary and non-redundant roles in dictating polarity to control ACD.

These experiments reveal a new paradigm by which Notch controls cell fate. In developing T cells, Notch plays a key role in dictating its own asymmetry during cell division and influencing asymmetry of other fate determinants. This work focused on developing T cells, but given the influence of Notch on Numb asymmetry in neural precursor cells, this might be a more general paradigm.

## Acknowledgments

We thank Juan Carlos Zúñiga-Pflücker (University of Toronto, Sunnybrook Health Sciences Centre, Toronto, Canada), for OP9 stromal cell lines. This work was performed in part at the Biointerface Engineering Hub @ Swinburne, part of the Victorian node of the Australian National Fabrication Facility (ANFF), a company established under the National Collaborative Research Infrastructure Strategy to provide nano-and micro-fabrication facilities for Australia’s researchers. We thank Louise Cheng (Peter MacCallum Cancer Centre) and Helena Richardson (La Trobe University) for comments on the manuscript.

## Methods

### Primary developing T cell co-culture

Mouse hematopoietic stem cells (isolated from C57BL/6 fetal liver at E14.5) were seeded onto OP9-DL1 stromal cells (received from Juan-Carlos Zúñiga-Pflücker, University of Toronto, Sunnybrook Health Sciences Centre, Toronto, Ontario, Canada) at a 1:1 ratio in 6 well plates (2 × 10^5^) in Minimal Essential Medium Alpha Modification supplemented with foetal calf serum (10% v/v), glutamine (1 mM), β-mercaptoethanol (50 µM), sodium pyruvate (1 nM), HEPES (10 mM), penicillin/streptomycin (100 ng/mL), mouse interleukin 7 (IL-7, 1 ng/mL) and mouse FMS-like tyrosine kinase 3 (Flt-3, 5 ng/mL). Hematopoietic cells were harvested via forceful pipetting and co-cultured on fresh OP9-DL1 stromal cells every 3–8 days. All mice were maintained in a specific pathogen-free environment with food and water freely available. All experiments on mice were performed in accordance with the Animal Experimentation Ethics Committee of the Peter MacCallum Cancer Centre.

### Retroviral transduction

Phoenix E cells (provided by Garry Nolan, Stanford University, Stanford, CA) were maintained at 37°C and at 10% CO_2_ in Dulbecco’s Minimal Essential Medium supplemented with foetal calf serum (10% v/v), L-glutamine (1 mM) and penicillin/streptomycin (100 ng/mL). Calcium phosphate transfection was performed on Phoenix E cells with 10-20 μg of the following pMSCV retroviral constructs: Cherry, GFP, GFP-Numb, Cherry-Numb, Cherry-α-Adaptin, and GFP-α-Adaptin in 10 cm dishes (Corning). Viral supernatant was harvested 48 h after transfection and added to 6 well plates that had been precoated with 15 μg/ml RetroNectin (Takara Bio Inc.) and blocked with 2% BSA. After addition of the viral supernatant, plates were spun at 2,000 *g* for 1 h and incubated for 1 h at 37°C. 5 × 10^5^ hematopoietic cells (day 4 co-culture) were added, and plates were spun for 1 h at 1,200 *g*.

### Flow cytometry

Developing T cell subsets were purified by staining for the cell surface markers CD44 and CD25 to discriminate between DN 1-4, CD28 to discriminate between DN3a and DN3b, and CD4 and CD8 to discriminate between the DN and DP populations. Hematopoietic cells were harvested from OP9-DL1 cocultures by forceful pipetting and stained on ice for one hour with the following antibodies: biotin-CD4 (1:2000; BD Pharmingen, 553045), biotin CD8a (1:2000; BD Pharmingen, 553029), PerCP/Cy5.5 CD44 (1:300; Biolegend 103032), APC-eFluor 780 CD25 (1:300; eBioscience, 47-0251-82) and Pe/Cy7 CD28 (1:200; Biolegend, 122014). Subsequently, the DN3a cells were rinsed and stained with BV421 Streptavidin (1:600, BD Biosciences, 563259) for one hour on ice prior to sorting. DN3a cells were isolated on a FACS (FACS Aria III; BD Biosciences) based on surface expression (CD28^lo^/CD25^+^/CD44^lo^/CD4^−^/CD8^−^). Retrovirally transduced cells were sorted 72 hours after transduction and also sorted on the basis of GFP and Cherry fluorescence. For the analysis of T cell differentiation, cells were stained with Pacific Blue CD4 (1:600; BD Pharmingen, 558107), APC-eFluor 780 CD8a (1:300; eBioscience, 47-0081-82), PerCP/Cy5.5 CD44 (1:300; Biolegend 103032), APC CD25 (1:300; eBioscience, 17-0251-82), Pe/Cy7 CD28 (1:200; Biolegend, 122014) and Sytox green viability stain (30 nM; Molecular Probes, S34860).

### Substrate functionalisation

Functionalised surfaces were used to individually present ligands to the developing T cells. Protein A (PA) was used to couple the Fc linked ligands to the surface to ensure optimum orientation of the ligand (Makaraviciute and Ramanaviciene, 2013, Song et al., 2012, Toda et al., 2011). Glass coverslips, silicon dioxide beads (5 µm), PDMS cell paddocks or 24 well plates were coated with protein A (235 nM, 1 hour), rinsed with PBS and further functionalised, if required, with Fc-DL1 (86 nM; R&D Systems, 5026-DL), Fc-DL4 (86 nM; BioVision, P1163), Fc-VCAM-1 (19 nM; R&D Systems, 643VM) or Fc-DL4 and Fc-VCAM-1 (86 nM + 19 nM) for 30 minutes. The substrates were then washed in PBS and media before cell deposition. For live cell imaging PDMS cell paddocks (120 µm x 120 µm) (Day et al., 2009) were rendered hydrophilic by exposure to air plasma at 1.5 × 10^-2^ mbar for 10 minutes and placed into an 8 well chamber slide (Ibidi) prior to protein functionalisation.

### Proliferation and development of DN3 cells on the functionalised surfaces

For the analysis of the ability of functionalised surfaces to support developing T cell proliferation and differentiation, 24 well plates were coated with protein A, protein A plus Fc-linked proteins or OP9-DL1 stromal cells. 1 × 10^5^ DN3 cells were seeded into the wells and assessed at 1 and 5 days by flow cytometry.

### Immunofluorescence and fixed image acquisition by confocal microscopy

2 × 10^4^ DN3a cells were added to 8 well chamber slide (Thermo Fisher Scientific) precoated with protein A, protein A plus Fc-linked proteins or OP9-DL1 stromal cells and cultured for 2 or 15 hours. For stromal cell-free culture cells were cultured in the presence of Flt-3 (5 ng/mL), CXCL12 (10 nM) and IL-7 (10 ng/ml) (Janas et al., 2010). Cells were then fixed, permeabilized, and labelled with primary antibodies for α-tubulin (1:800; Rocklands, 600-401-880 or 200-301-880) and test proteins, Notch1 (1:200; Abcam, ab27526), α-Adaptin (1:200; Abcam, Ab2730), Numb (1:200; Abcam, Ab4147), CXCR4 (1:200; BD Pharmingen, 551968), CD25 (1:1000; BD Pharmingen, 553070), and the appropriate secondary antibodies and then mounted with Prolong gold antifade (Molecular Probes). The slides were examined using an Olympus FV1000 microscope (Olympus) and 40x objective lens (0.95 NA Oil Plan Apochromat) or Olympus FV3000 microscope (Olympus) and 60x objective lens (1.35 NA Oil Plan Apochromat). 3D images of the cells were acquired with a z distance of 0.5 μm and maximum intensity projections of z sections created using ImageJ. For quantifications, the investigator was blind to the sample allocation at the moment of scoring. For the analysis of polarisation during interphase α-tubulin staining was used to identify cells with the MTOC at the cell-cell / cell-substrate interface and the cell was cored depending on whether the protein of interest was diffuse or enriched at the MTOC (**Fig. S8A**). To determine polarisation during division mitotic cells were identified by the presence of a mitotic spindle and the cell division was assigned as symmetric if the fluorescence of the test protein was evenly distributed between the two daughter cells and asymmetric if the fluorescence was greater in one of the daughter cells than the other daughter cell (**Fig. S8B**). The co-inheritance of Notch with other fate determineants (specifically Numb, α-Adaptin and CXCR4) was assessed by selecting dividing cells which were asymmetric for both proteins for analysis and scoring it as co-inherited if both proteins were concentrated within the same nascent daughter cell.

### Live cell image acquisition and analysis

For live cell imaging, 2 × 10^4^ DN3a cells were added to 8 well chamber slide (Ibidi) with the PDMS cell paddocks precoated with protein A, Fc-linked proteins or OP9-DL1 stromal cells. For stromal cell-free culture cells were cultured in the presence of Flt-3 (5 ng/mL), CXCL12 (10 nM) and IL-7 (10 ng/mL) (Janas et al., 2010). Images were captured on a spinning disc confocal system fitted with an inverted microscope and temperature controlled chamber maintained at 37°C and 5% CO_2_. Images are acquired using a 40x air objective (0.95 NA) and multiple stage positions are recorded every 3 min for 20 hours, with 8-slice z-stacks of 1 µm thickness. As described in (Charnley and Russell, 2017, Oliaro et al., 2010, Pham et al., 2015, Shimoni et al., 2014), the polarisation of fluorescent markers in the nascent daughter cells was quantified by measuring the total fluorescence intensity of each daughter cell and applying a “Polarization Ratio” (PR) equation: PR = (∑H1 - ∑H2)/ (∑H1 + ∑H2); where the difference in total intensity between Daughter 1 (∑H1) and Daughter 2 (∑H2) is divided by the sum of intensities in both Daughter 1 (H1) and Daughter 2 (H2). PR of 0.17 was used as a cutoff to designate cells as undergoing ACD (PR > 0.17) or SCD (PR < 0.17). On the functionalised surfaces, only cells that had not interacted with another cell for at least 30 minutes prior to division were selected for analysis. This enabled the isolation of the effect of the immobilised protein on ACD. To compare the distribution of PR α-Adaptin values between the different cell culture platforms, outliers (which were identified by their high PR diffuse values and consisted of less than 10% of dividing cells) were removed (Charnley and Russell, 2017) and the data was re-plotted as violin plots using the kernel density function.

To assess the role of the orientation of the polarity cue, 5 µm silicon dioxide beads were coated with PA or PA + Fc-DL1 using the same protocol outlined for the 2D substrates. The orientation of division relative to the polarity cue was determined for dividing DN3a cells in contact with functionalised beads and OP9-DL1 stromal cells. For DN3a cells in contact with OP9-DL1 stromal cells, the division angle was determined by measuring the angle between the interface between the DN3a cell and OP9-DL1 stromal cell and the long axis of the dividing cell (**Fig. S8C**). For the functionalised beads, a line was drawn between the centre of mass of the bead and the centre of mass of the dividing cell. The division angle was defined as the angle orthogonal to the angle between this line and the long axis of the dividing cell (**Fig. S8D**).

### Notch and CXCR4 inhibition protocol

To disrupt Notch signalling we used the γ-secretase inhibitor, dibenzazepine (DBZ; 0.05 µM) (van Es et al., 2005). DBZ prevents the cleavage of Notch by γ-secretase, and so prevents the translocation of the cytoplasmic portion of Notch to the nucleus and therefore inhibits Notch signalling. To analyse the effect of Notch signalling on proliferation and differentiation, purified DN3 cells were co-cultured on OP9-DL1 stromal cells in the presence or absence of DBZ and assessed by flow cytometry. To determine the effect of Notch and CXCR4 inhibition on ACD, purified DN3a cells were cultured on OP9-DL1 stromal cells or Fc-DL1 functionalised surfaces in the presence of AMD3100 (2 μg/ml, Sigma-Aldrich) or DBZ (0.05 µM, Calbiochem) for approximately 30 mins prior to the start of the time lapse imaging or for 15 hours prior to fixing for immunofluorescence analysis.

### Statistical Analysis

All data was collected from 2 to 8 independent experiments. For scatter and bar plots data is shown as mean ± SEM, median value is indicated on the violin plots. The following tests were used for the statistical analysis: Student’s unpaired two-way t-test with unequal variance was used to determine the difference between mean values for the scatter plots and calculated using Microsoft Office Excel. For the PR scatter plots from the time lapse microscopy experiments paired *t* tests were performed in Graphpad Prism between control (diffuse protein) and test protein PR values. The violin plots were analysed for differences in the distribution of the data using Two-sample Kolmogorov-Smirnov (KS) test (Young, 1977) using Graphpad Prism software. Level of statistical significance is indicated as: *p < 0.05, **p < 0.01, ***p < 0.001.

---

**Figure S1.**
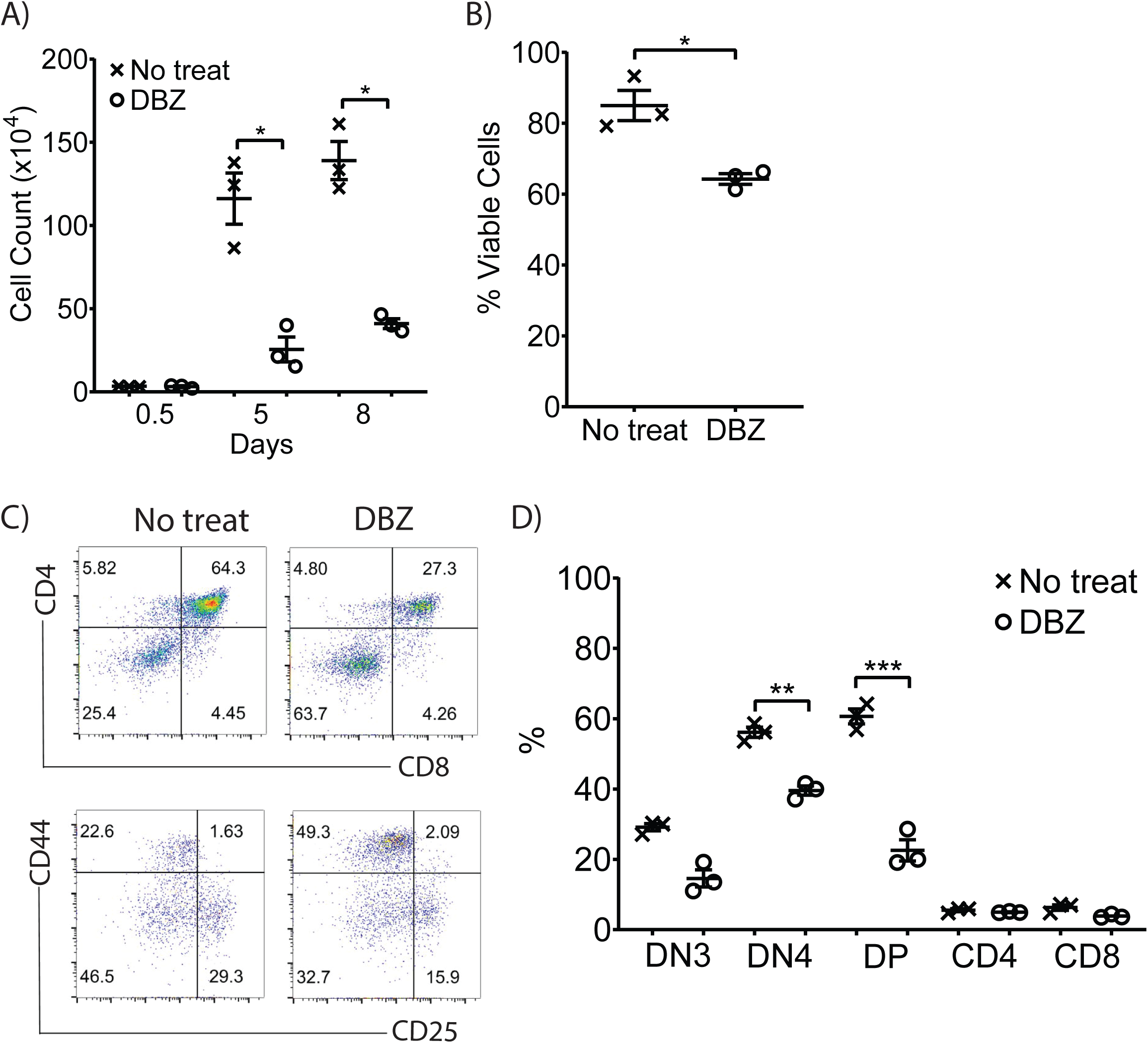
Notch signalling is important for downstream DN3 proliferation, survival and differentiation. DN3 cells were seeded on OP9-DL1 cells in the absence (no treat) or presence of γ-secretase inhibitor (DBZ; 0.05 µM) for up to 8 days and assessed using flow cytometry. **(A)** The inclusion of DBZ resulted in a reduction in the number of cells after 5 and 8 days of culture and **(B)** viability. **(C)** Flow cytometry plots from one representative experiment. **(D)** Treatment with DBZ reduced differentiation, evidenced by a decrease in DN4s and DPs after 8 days of culture. *n* = 3 independent experiments, all data are represented as mean ±SEM; *p < 0.05, **p < 0.01, ***p < 0.001 (unpaired t test).

**Figure S2.**
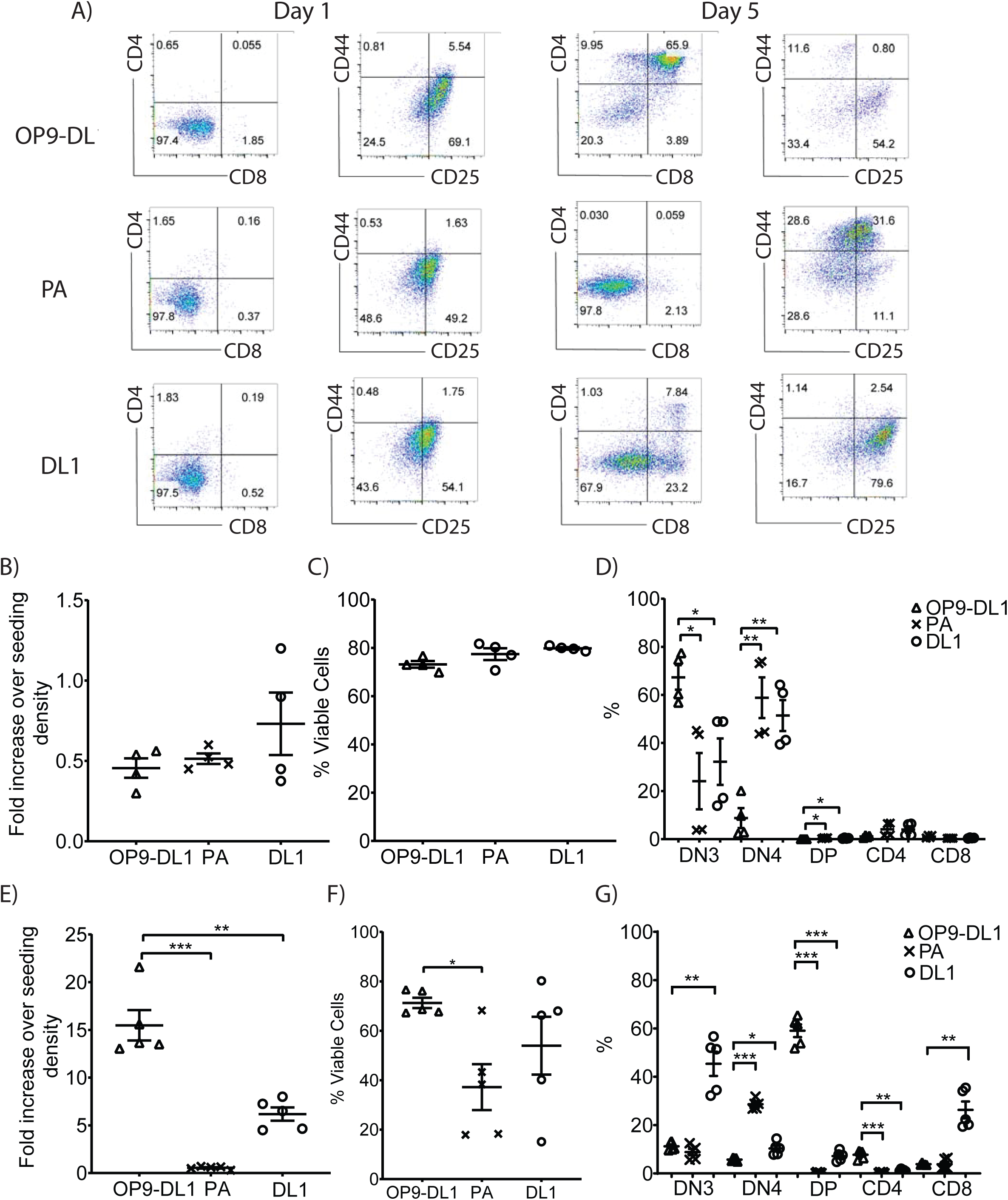
DL1 functionalised surfaces support DN3 proliferation and differentiation. DN3 cells were cultured onto surfaces pre-coated with PA, Fc-DL1 (DL1) or OP9-DL1 stromal cells and analysed with flow cytometry. **(A)** Flow cytometry plots from one representative experiment. Cells were cultured on the three surfaces for 1 day **(B, C, D)** or 5 days **(E, F, G)** and assessed for expansion, viability, and differentiation. Surfaces functionalised with Fc-DL1 supported DN3 development, although the expansion and development was reduced relative to culture on OP9-DL1 cells. Conversely, PA alone was not able to support thymocyte culture, as indicated by a substantial reduction in proliferation, increased cell death and a failure to develop past the DN stage. *n* = 5 independent experiments, all data are represented as mean ±SEM; *p < 0.05, **p < 0.01, ***p < 0.001 (unpaired t test).

**Figure S3.**
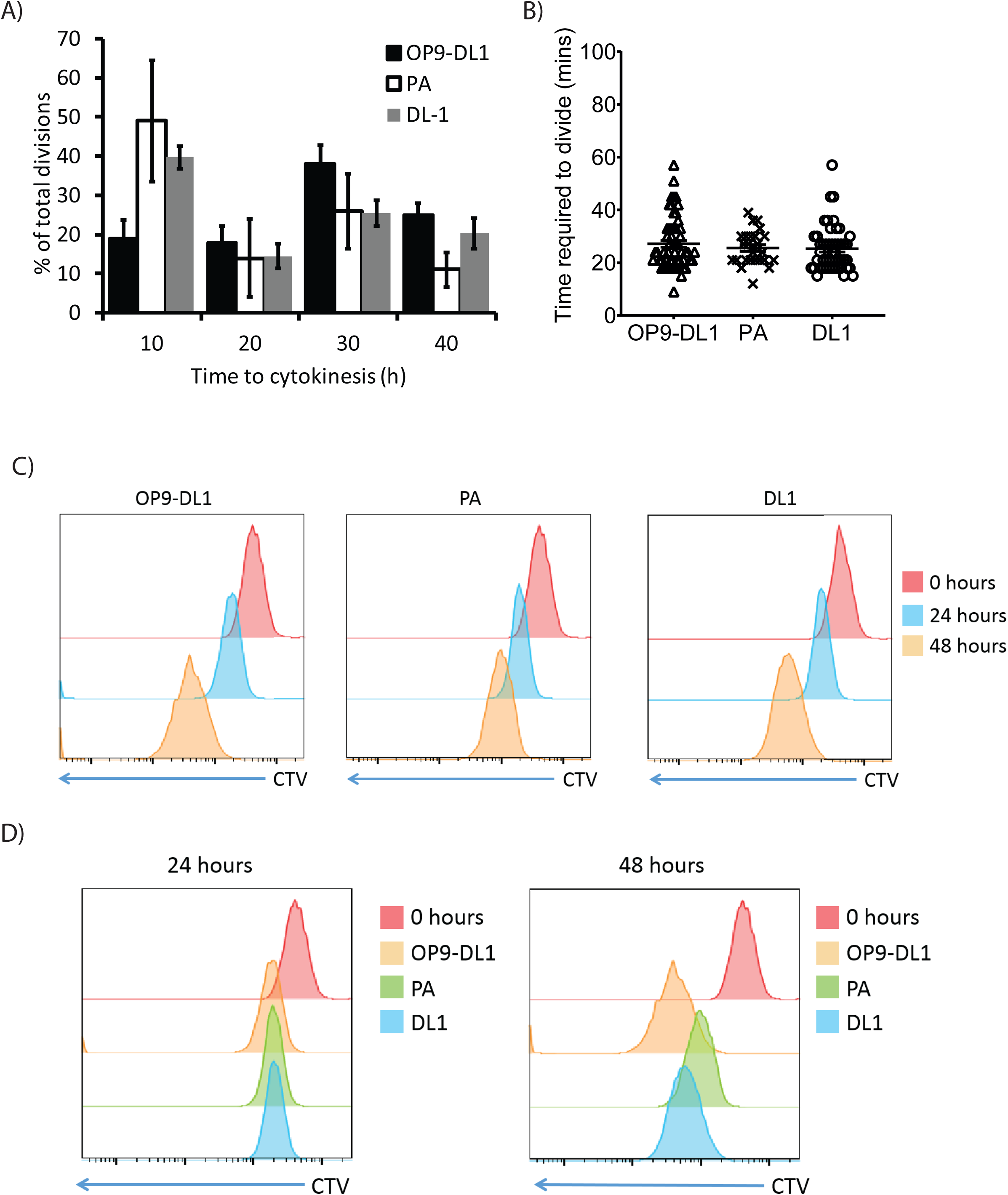
Functionalised surfaces support the division of DN3a cells. **(A)** DN3a cells were cultured on OP9-DL1, PA and Fc-DL1 (DL1) for up to 40 hours. The time of divisions (defined as time that cytokinesis occurred) were binned into 10 hours intervals and expressed relative to the total number of divisions on that surface. Cells divided throughout the movies, although there was a decrease in the number of divisions on the PA surfaces at later timepoints. **(B)** The time required for the DN3a cells to divide (defined as nuclear envelope breakdown to cytokinesis) was not affected by the cell culture platform. **(C)** Cell Trace Violet (CTV) data for DN3a cells cultured on the functionalised surfaces and OP9-DL1 stromal cells revealed that thymocytes divided on all of the cell culture platforms. **(D)** However, after 48 hours division was reduced on the PA surfaces and to a lesser extent on the Fc-DL1 functionalised surfaces. *n* = 2-7 independent experiments, with flow cytometry plots from one representative experiment shown. All data are represented as mean ±SEM.

**Figure S4.**
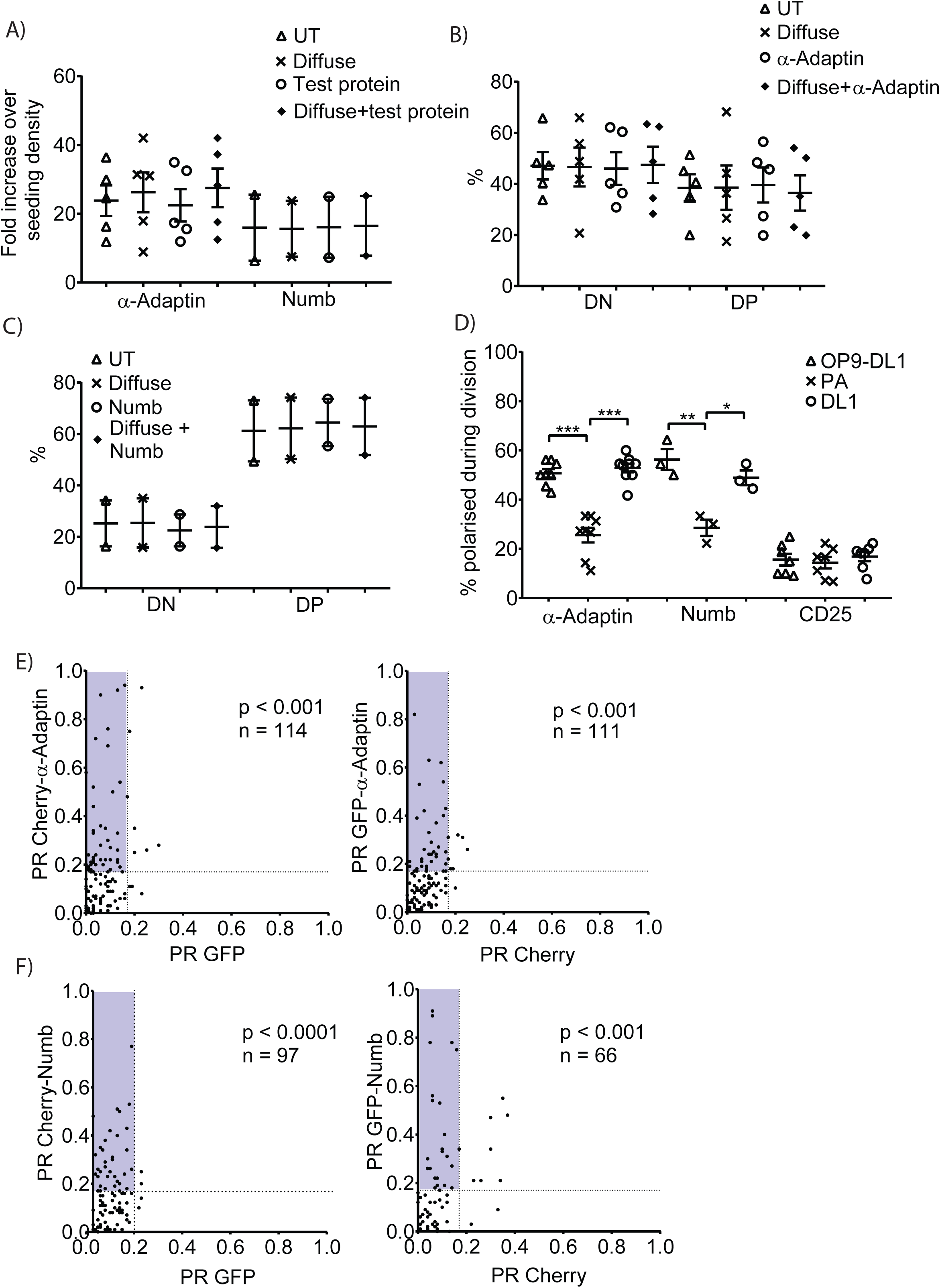
Overexpression of α-Adaptin or Numb does not affect proliferation, differentiation or polarisation during division. DN3a cells expressing a diffuse protein (GFP or cherry) and α-Adaptin or Numb were cultured on OP9-DL1 stromal cells for 5 days and assessed using flow cytometry. **(A)** Transduction with either α-Adaptin or Numb had no effect on proliferation. The overexpression of **(B)** α-Adaptin or **(C)** Numb also had little effect on differentiation. **(D)** DN3a cells expressing a diffuse protein (GFP or cherry) and α-Adaptin were cultured onto surfaces pre-coated with PA or Fc-DL1 (DL1) for 15 hours. Dividing cells were identified from α-tubulin staining and scored as polarised or symmetric. The proportion of cells that were polarised was comparable to the proportion observed when assessed by antibody labelling of endogenous protein (compare with Fig. 4). Total number of divisions analysed: OP9-DL1 = 97, 52 and 45; PA = 99, 59 and 40; Fc-DL1 = 107, 55 and 52 for GFP, α-Adaptin and Numb, respectively. **(E)** PR scatter plots for dividing DN3a cells transduced with either GFP and cherry-α-Adaptin or GFP-α-Adaptin and cherry and cultured in contact with OP9-DL1 stromal cells. **(F)** PR scatter plots for dividing DN3a cells transduced with either GFP and cherry-Numb or GFP-Numb and cherry and cultured in contact with OP9-DL1 stromal cells. PR was calculated as the difference in fluorescence between the two halves divided by the sum of fluorescence. The distribution of PR values was similar regardless of whether the test protein was coupled to GFP or cherry. Total number of divisions analysed and p for the comparison of diffuse *versus* test protein PR values (paired t test) are shown. The blue shaded region indicates divisions that were assigned as asymmetric when the 0.17 cut-off was applied. *n* = 2-9 independent experiments, all data are represented as mean ± SEM; *p < 0.05, **p < 0.01, ***p < 0.001 (unpaired t test).

**Figure S5.**
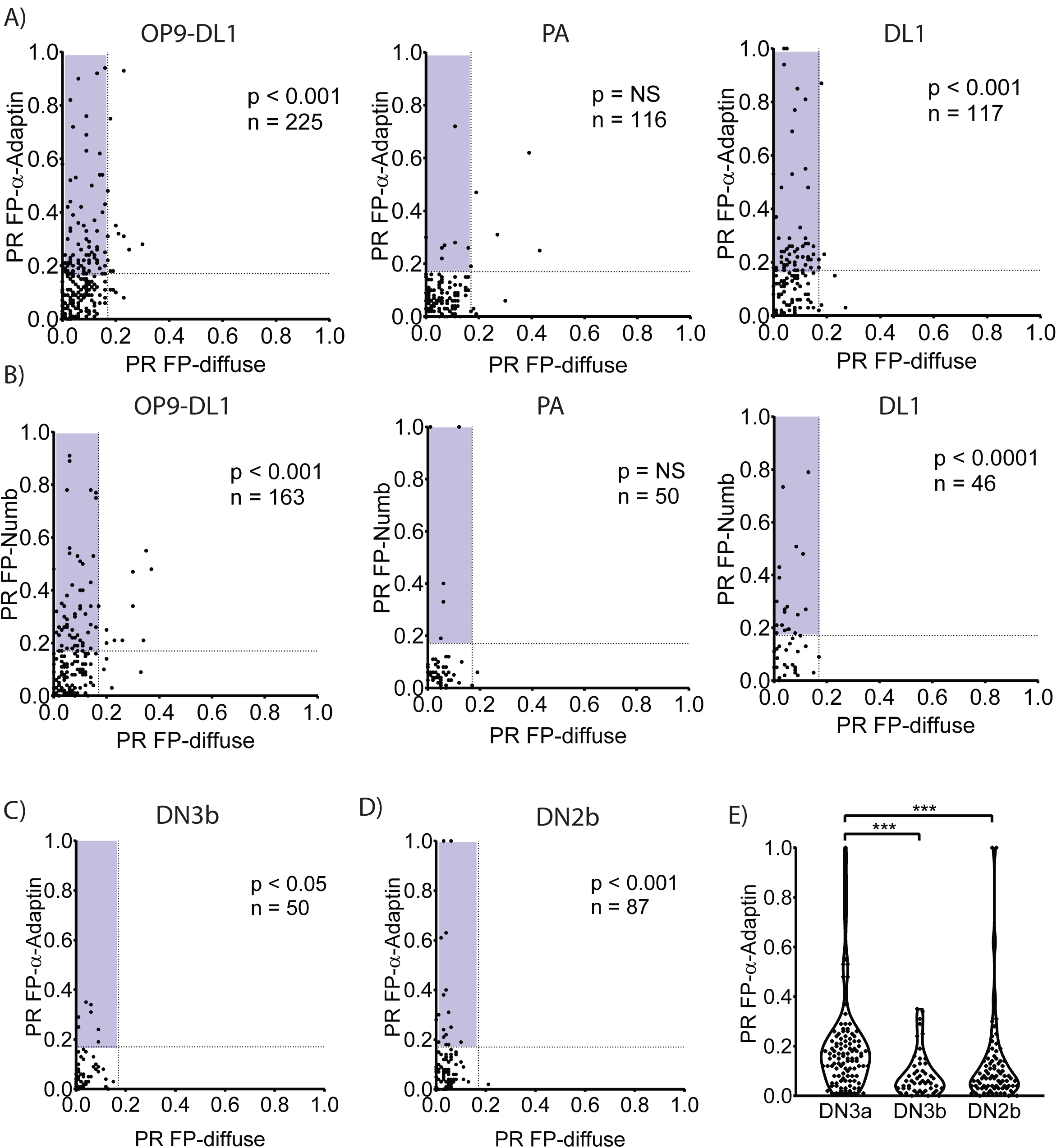
DN3a cells undergo polarised divisions on functionalised surfaces. PR scatter plots for dividing DN3a cells expressing a diffuse protein (GFP or cherry) and **(A)** α-Adaptin or **(B)** Numb cultured in contact with OP9-DL1 stromal cells, PA or Fc-DL1. Culturing DN3a cells on OP9-DL1 stromal cells and Fc-DL1 increased the proportion of dividing cells with high PR values for α-Adaptin and Numb. Polarised divisions were stage specific as indicated by a lack of polarised divisions in **(C)** DN3b cells and **(D)** DN2b cells. Total number of divisions analysed and p for the comparison of diffuse *versus* test protein PR values (paired t test) are shown. The blue shaded region indicates divisions that were assigned as asymmetric when the 0.17 cut-off was applied. **(E)** Violin plot of DN3b and DN2b *versus* DN3a. Number of divisions analysed: DN3a = 109, DN3b = 50 and DN2b = 86; *n* = 2-4 independent experiments; magenta line on violin plots indicates median value; ***p < 0.001 (KS test).

**Figure S6.**
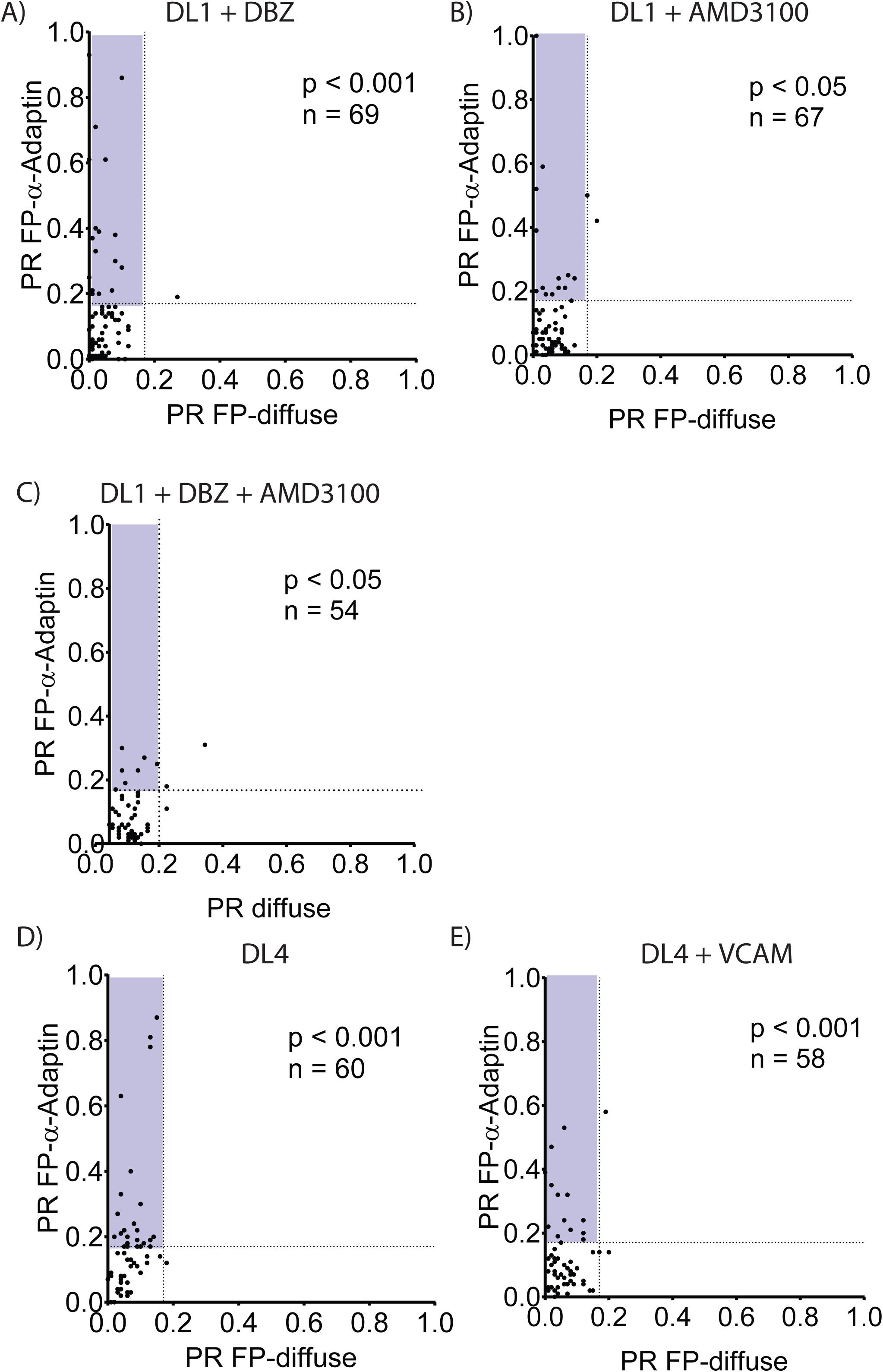
Different polarity cues can promote or inhibit ACD. DN3a cells expressing a diffuse protein (GFP or cherry) and α-Adaptin were cultured onto surfaces pre-coated with PA, Fc-DL1 (DL1), Fc-DL4 (DL4) or Fc-DL4 + VCAM-1 (DL4 + VCAM) and imaged using time-lapse microscopy for 20 hours. The addition of **(A)** DBZ or **(B)** AMD3100, to inhibit Notch or CXCR4 signalling, respectively, partially abolished ACD. **(C)** The inclusion of both inhibitors further reduced the level of ACD. **(D)** Fc-DL4 coated surfaces triggered ACD. **(E)** The inclusion of VCAM-1 lead to a reduction in ACD. Blue shaded regions in PR scatter plots indicates divisions that were assigned as asymmetric using the 0.17 cut-off, total number of divisions analysed and p for the comparison of diffuse *versus* test protein PR values (paired t test) are shown. n = 2-4 independent experiments.

**Figure S7.**
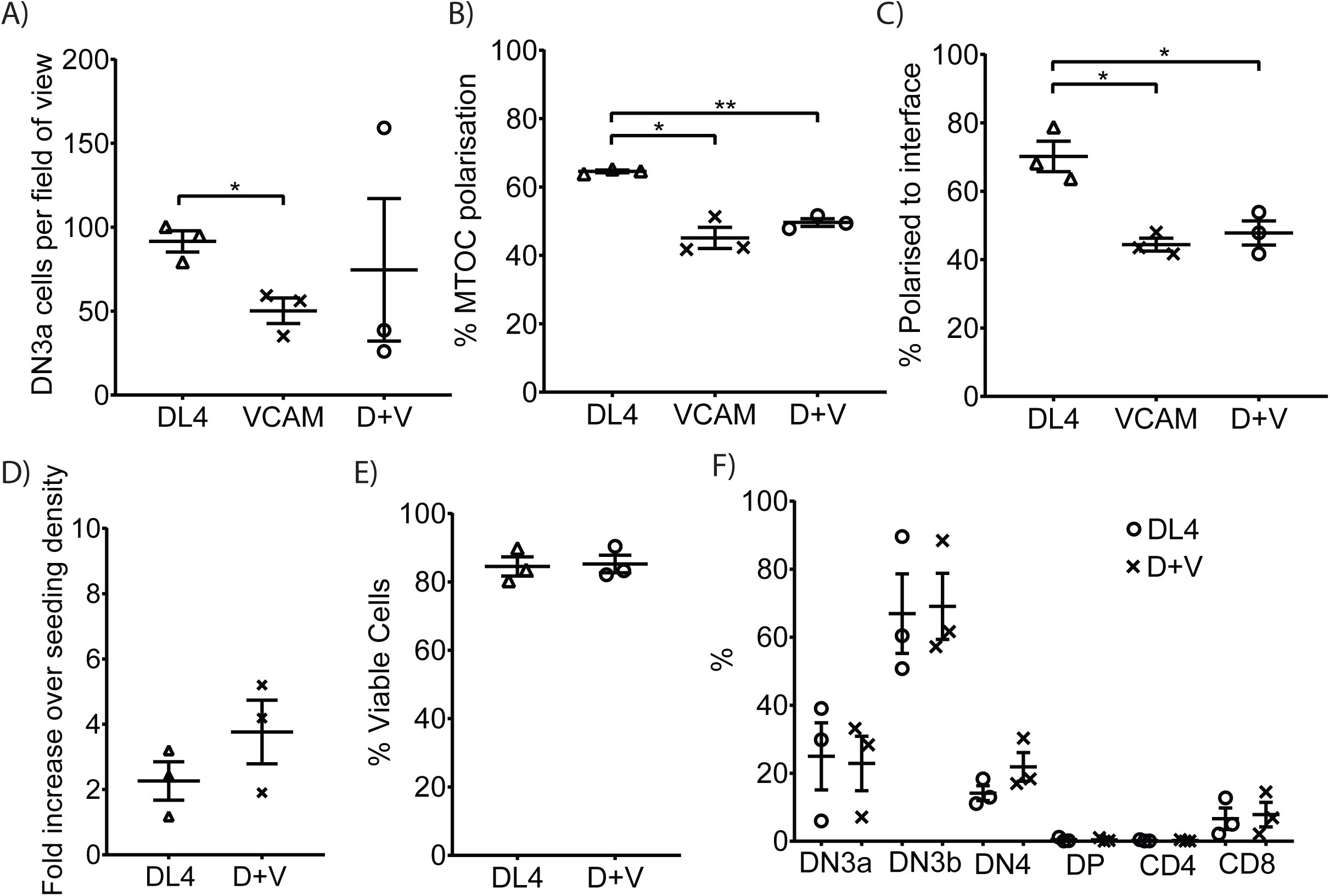
The addition of Fc-VCAM-1 to Fc-DL4 coated surfaces reduces polarisation during interphase. Sorted DN3a cells were cultured onto surfaces pre-coated with Fc-DL4 (DL4), Fc-VCAM-1 (VCAM) or Fc-DL4 + Fc-VCAM-1 (D+V) for 15 hours before fixing and staining with α-tubulin and α-Adaptin. **(A)** Thymocyte attachment was reduced on surfaces coated by VCAM-1 alone. **(B)** MTOC recruitment and **(C)** α-Adaptin polarisation to the cell-substrate interface was reduced in the presence of Fc-VCAM-1. Total number of divisions analysed: DL4 = 113, VCAM-1 = 59, DL4 + VCAM-1 = 61. DN3a cells were cultured onto surfaces pre-coated with Fc-DL4 (DL4) or Fc-DL4 + Fc-VCAM-1 (D+V) for 5 days before harvesting and analysis with flow cytometry. The inclusion of Fc-VCAM-1 had no effect on **(D)** cell number, **(E)** viability or **(F)** differentiation. All data are represented as mean ± SEM; *n* = 3 independent experiments, *p < 0.05, **p < 0.01 (unpaired t test).

**Figure S8.**
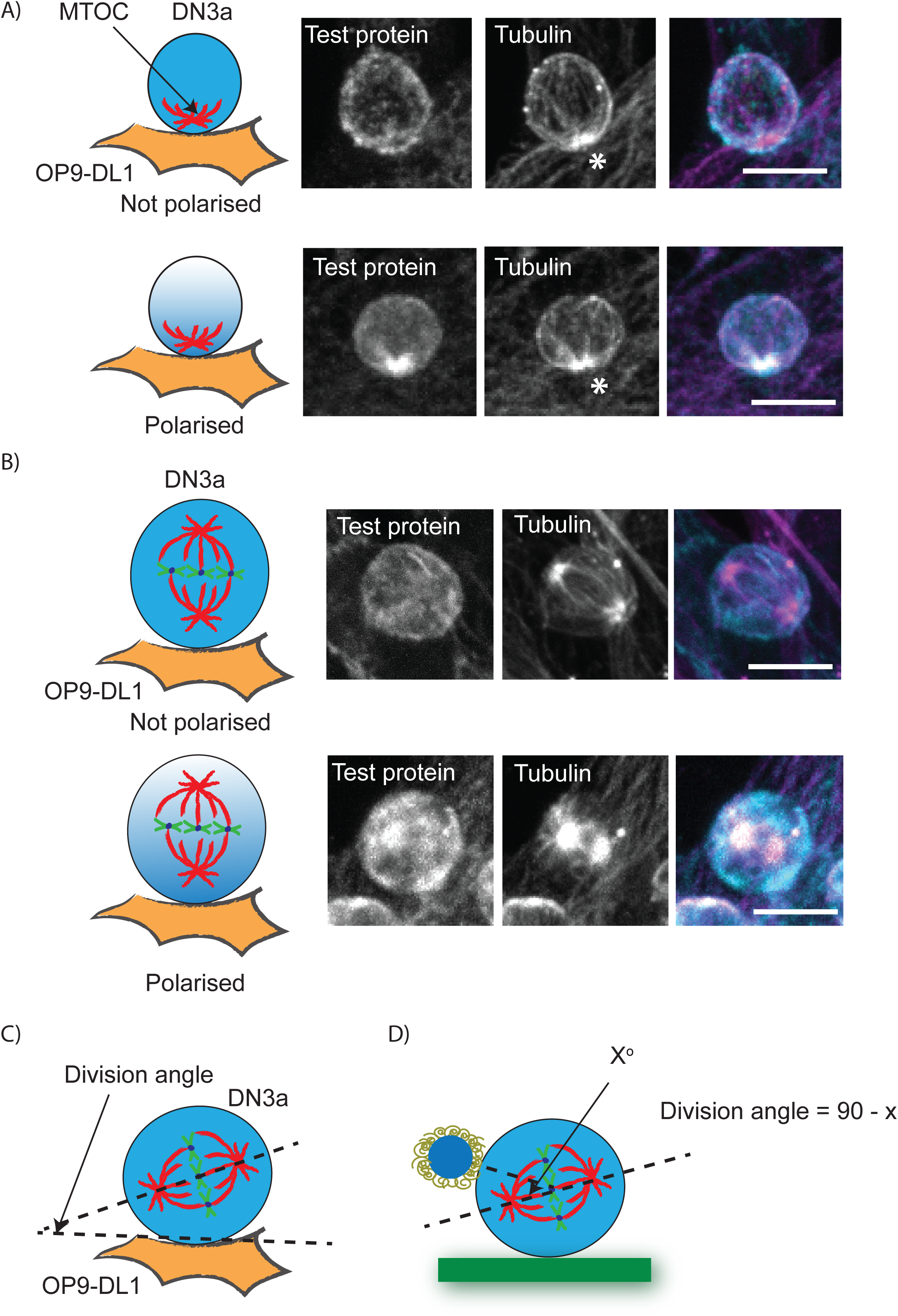
Analysis of polarisation during interphase and division in DN3a cells. **(A)** To determine polarity during interphase, cells in which the MTOC was recruited to the stromal interface were selected for analysis and the protein of interest was assigned as diffuse or polarised, where polarisation was defined as a clear enrichment of fluorescence at the interface. * indicates the interface between the DN3a and OP9-DL1 stromal cells. Schematic of analysis is shown on the left and representative projected z stack image is shown on the left. Bar = 10 µm. **(B)** To determine polarisation in dividing cells dividing cells were identified from α-tubulin staining and scored as undergoing SCD if the protein of interest was symmetrically distributed between the two daughter cells or ACD if the protein of interest was more concentrated in one of the daughter cells. Schematic of analysis is shown on the left and representative projected z stack image is shown on the left. Bar = 10 µm. Schematic to demonstrate the measurement of the orientation of mitotic spindle relative to the polarity cue. **(C)** For DN3a cells cultured in contact with OP9-DL1 stromal cell the division angle was defined as the angle between the the DN3a cell and OP9-DL1 stromal cell interface and the long axis of the dividing cell. **(D)** For the functionalised beads, a line was drawn between the centre of mass of the bead and the centre of mass of the dividing cell. The division angle was defined as the angle orthogonal to the angle between this line and the long axis of the dividing cell.

